# Recurrent leukaemia stem cell states define the antigenic landscape of paediatric Acute Myeloid Leukaemia

**DOI:** 10.64898/2026.10.19.760044

**Authors:** Sina Kanannejad, Macarena Oporto Espuelas, Claudia D’Oria, Filippo Martignano, Ilaria Marina Michelozzi, Soumick Chatterjee, Edoardo Giacopuzzi, Eduardo Gomez Castaneda, Aurora Casiraghi, Theodoros Xenakis, George T. Hall, Efstratios Kirtsios, Cecilia Reyneri, Hannah Macpherson, Jack Bartram, Rebecca Thomas, Sergi Castellano, Persis J. Amrolia, Alice Giustacchini

**Author notes:** These authors contributed equally to this work (Sina Kanannejad, Macarena Oporto Espuelas).

## Abstract

Paediatric acute myeloid leukaemia (pAML) remains a leading cause of childhood cancer mortality, with disease heterogeneity representing a major obstacle to effective immunotherapy. To enable rational immunotherapy design accounting for such heterogeneity, we combined single-cell transcriptomics and surface proteomics (CITE-seq). Using an integrated framework for malignant cell identification and unbiased gene program identification, we resolved three conserved leukaemia stem cell (LSC) subtypes, primitive (P-LSC), monocytic (M-LSC), and erythroid-like (E-LSC), that frequently coexist within individual patients in varying proportions and represent distinct stem-cell states with characteristic immunophenotypes and transcriptional programmes. Quantitative surface antigen profiling revealed that no single target achieved full LSC coverage. CLEC12A, an antigen currently being evaluated clinically, was expressed across P-LSC and M-LSC but was largely absent from E-LSC. In contrast, CD84 provided complementary coverage in the P and M groups and served as the primary target in the E-LSC population. The combination of CLEC12A and CD84 provided broad coverage across all three LSC subtypes, while sparing healthy haematopoietic stem cells. Validation across independent single-cell and bulk cohorts confirmed the robustness of this combination. These findings establish a paediatric-specific framework for antigen prioritisation in AML, identify CLEC12A/CD84 as a robust dual-targeting strategy across heterogeneous LSC states, and provide a blueprint for the rational design of next-generation CAR T-cell combinations.

## Introduction

Paediatric acute myeloid leukaemia (pAML) is a rare but aggressive haematological malignancy that remains a leading cause of cancer-related mortality in children. Relapse is the primary cause of treatment failure: children who relapse have long-term survival of 40%, with no curative therapy after allogeneic haematopoietic stem cell transplantation [1]. Outcomes have not significantly improved over the last two decades, and novel therapies are therefore urgently needed[2].

AML is a molecularly heterogeneous group of diseases with a complex mutational landscape. Recurrent cytogenetic and molecular abnormalities have prognostic utility when classifying patients at high risk of relapse and have been applied to risk-adapted front-line therapy. Clinically, paediatric AML is classified according to the same World Health Organization (WHO) and International Consensus Classification (ICC) diagnostic frameworks and risk stratification guidelines used in adult AML[3–5]. In children, fusion-driven events such as KMT2A rearrangements, RUNX1::RUNX1T1, CBFB::MYH11, NUP98 rearrangements, and GLIS fusions are common and constitute major molecular subgroups of AML[6, 7]. Recent advances in pAML genomic profiling have contributed to refining these categories, leading to new molecular classes with distinct transcriptional profiles that may enable a more accurate prognostic framework[7].

Interestingly, while pAML transcriptional profiles often align with molecularly defined classes, they can also help highlight phenotypic diversity. For example, KMT2A- and NUP98-rearranged leukaemias comprise patients with transcriptional states resembling both myelomonocytic (M4/M5) and erythroid/megakaryocytic (M6/M7) lineages. Single-cell RNA sequencing has further illustrated such immunophenotypic heterogeneity by mapping pAML cells onto a normal haematopoietic reference, revealing patient-specific differentiation blocks at the level of discrete cell populations[8]. These analyses have identified recurrent AML differentiation programmes, ranging from primitive to lineage-biased, with immunophenotypes that can diverge even among patients sharing the same driver events and from diagnosis to relapse, with important implications for therapeutic targeting[8, 9].

Immunophenotypic diversity is further complicated by intra-clone functional heterogeneity, according to which malignant cells can differ in their self-renewal ability, proliferation, and therefore therapy resistance. In line with this, leukaemia stem cells (LSCs) are believed to play a central role in disease persistence and relapse in AML, and high LSC transcriptional scores are associated with poor prognosis in both adult and paediatric AML[10–14]. Recent single-cell studies have further refined our understanding of LSC and blast biology, identifying functionally distinct LSC sub-types, including primitive and monocytic populations, with divergent drug sensitivity profiles, as well as distinct transcriptional programmes [15–20].

Altogether, the genomic, immunophenotypic, and functional complexity of pAML underscores the need for therapeutic strategies that can eradicate the full spectrum of malignant cell states, spanning diverse immunophenotypic and functional properties, particularly stem-like ones.

To this end, Chimeric Antigen Receptor (CAR) T cell therapy is agnostic to transcriptional or functional cell states and can, in principle, eliminate any cell expressing the surface antigens they are programmed against. However, translating the success of CAR T cell therapy in B-lineage haematological malignancies to AML has been hindered by the lack of AML-specific targets. Current clinical efforts in adult and paediatric AML have focused on common myeloid antigens, such as CD33, CD123 and CLEC12A alone or in combination, with early-phase trials reporting modest response rates, frequent early relapse, and limited CAR T cell persistence across targets[21, 22]. Clinical experience to date suggests that therapeutic efficacy is constrained by the lack of AML-specific antigens, resulting in on-target/off-tumour myeloablation, and by antigen heterogeneity across AML cell states, particularly at the level of LSCs, which allows residual disease to escape single-antigen targeting.

These challenges have motivated the development of alternative therapeutic strategies, including cooperative and dual-antigen CAR designs, as well as gene-edited haematopoietic grafts engineered to tolerate targeting of shared myeloid antigens[22, 23]. In parallel, approaches aimed at improving CAR T-cell persistence and activity within the AML microenvironment, such as IL-18-armoured CARs, are also entering clinical evaluation[24]. Despite their mechanistic differences, all of these approaches ultimately depend on a systematic, high-resolution antigen mapping across paediatric AML cell states.

Simultaneously resolving transcriptional programmes and surface protein expression at single-cell resolution is critical in this context, as transcript abundance alone does not reliably predict surface protein levels for candidate therapeutic targets. CITE-seq addresses this by enabling direct quantification of antigen expression across immunophenotypically and functionally defined malignant subpopulations, including rare LSC-like compartments. Pioneering works in adult AML[23, 25, 26], combining transcript and protein readouts, have contributed to profiling surface antigen expression across healthy and malignant immunophenotypic compartments, laying the basis for the rational design of CAR T cell target combinations. However, no study to date has systematically mapped surface antigens across cell states in paediatric AML.

Here, we combine single-cell transcriptomics with CITE-seq–based surface proteomics to profile a ntigen expression across malignant subpopulations in high-risk paediatric AML, enabling resolution on rare and clinically relevant compartments such as LSCs. We identify LSC-like transcriptional programs that are highly conserved across cytogenetic groups, despite differences in lineage bias. Building on this, we develop a framework to prioritise cell-state–resolved CAR T cell target identification, designed to eradicate the full spectrum of malignant cell states, spanning diverse immunophenotypic and functional properties, particularly stem-like ones. Within this framework, CLEC12A (CD371) and CD84 emerge as a lead target pair, together achieving broad coverage of the LSC-like population while sparing healthy HSCs. We validate this framework using both external single-cell datasets and bulk RNA-seq patient cohorts and confirm its ability to resolve surface antigen combinations that predict differential efficacy even within the same cytogenetic subtype. This approach provides a model for rational design of cell state–resolved immunotherapy target combinations that could be extended to other heterogeneous malignancies.

## 2 Results

### 2.1 CITE-seq profiling of a high-risk paediatric AML cohort

To systematically profile surface target expression across pAML transcriptional states, we generated a CITE-seq single-cell dataset of 27 bone marrow samples from 16 paediatric AML patients, as well as five healthy controls (two adult bone marrow, two cord blood, and one paediatric bone marrow). All patients were considered high risk, either based on adverse-risk cytogenetics or clinical relapse. Importantly, our cohort covered a wide range of cytogenetic subtypes, including relatively more common alterations (KMT2Ar, RUNX1::RUNX1T1, CBFB::MYH11, and NUP98r) as well as rarer fusions (GLISr, DEK::NUP214, and MNX1::ETV6), ensuring a broad coverage of molecular categories (Fig. 1a).

**Fig. 1.**
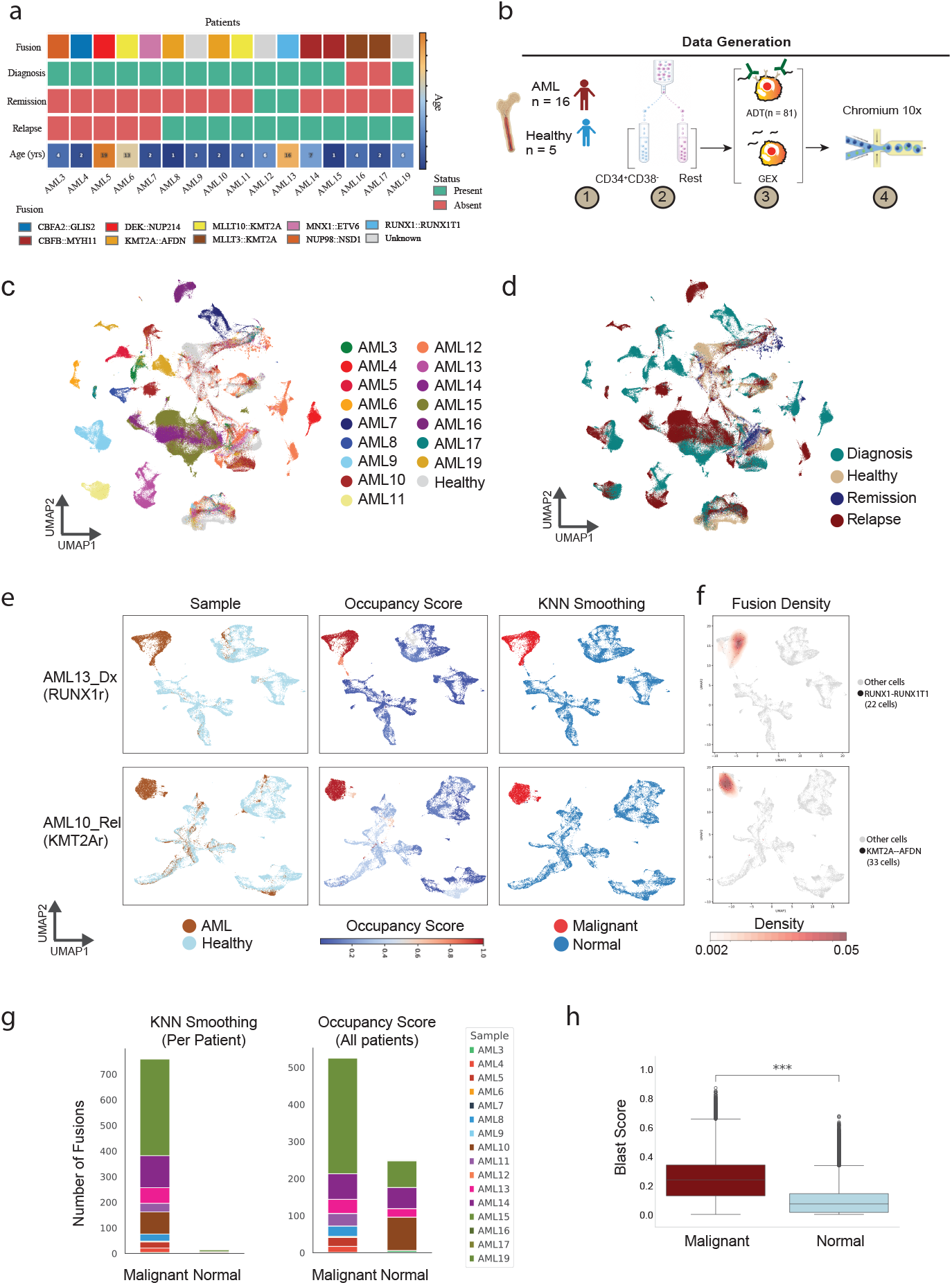
Paediatric AML cohort overview and malignant cell classification. **a**. Patient cohort overview. Single-cell CITE-seq was performed on 27 bone marrow samples from 16 high-risk paediatric AML patients and 5 healthy controls (2 adult bone marrow BM, 2 cord blood CB, and 1 paediatric bone marrow PBM), covering various common cytogenetic groups (Fusion). **b**. Experimental workflow. CD34^+^ flow cytometry enrichment was performed before CITE-seq library generation to enhance the capture of immature populations. **c**. UMAP of all single cells in our cohort, coloured by patient ID. **d**. UMAP of all single cells in our cohort, coloured by patient cytogenetic class. **e**,**f**. Malignant cell classification shown for two representative patients, AML13 diagnosis (dx) and AML10 relapse (rel). UMAPs show (left to right) sample of origin (AML vs. healthy donor), occupancy score, KNN-smoothed data, and fusion-specific cell density, highlighting cellular distribution and fusion-associated populations within individual patients. Healthy donor-derived cells in these plots include adult bone marrow and paediatric bone marrow. **g**. Comparison of malignant versus normal cell classification using occupancy scores with or without KNN smoothing. The number of patient-specific fusion events per cell is shown. In the occupancy score alone, cells are classified as malignant or normal using Leiden clustering with resolution = 2. **h**. Validation of KNN-smoothing classification by comparison with the blast gene expression score from Mumme et al.

To enrich for immature, stem-like populations often underrepresented in unsorted bone marrow mononuclear cells, our cohort underwent CD34-based enrichment (including CD34^+^CD38^−^ fractions) prior to single-cell capture, while retaining CD34^−^ cells to preserve full malignant and healthy immune microenvironment representation (Fig. 1b).

To capture both the transcriptional programs and surface antigen repertoires relevant to designing target combinations for CAR T cell strategies against pAML, we implemented CITE-seq with a custom panel of 81 oligo-tagged antibodies. These antigens were selected based on prior knowledge of LSC in AML and included targets under investigation in existing CAR T pipelines (Supplementary Table 3).

Following quality control, we retained 259,817 cells for downstream analysis, with a mean of 2,252 genes with associated RNA molecules per cell. For the protein modality, 69 of 81 antibody-derived tags (ADTs) were confidently detected (Methods), showing correlation with their corresponding RNA features and low correlation with isotype controls (Extended Data Fig. 1), with an average of 1,642 counts per cell. Importantly, to avoid filtering out viable cells with low RNA abundance, likely enriched for quiescent or LSC-like populations, we manually defined thresholds on transcript counts, gene counts, and mitochondrial content. This strategy ensured removal of low-quality cells while preserving high-quality ones, avoiding reliance on median-based filters or standard mitochondrial cutoffs, which have been shown to inadvertently discard relevant malignant clusters in tumours[27] (Extended Data Fig. 2). Clustering of all cells revealed the diversity of AML-derived populations, including patient-specific clusters from both diagnosis and relapse samples, recapitulating the fusion-induced heterogeneity observed among pAML patients (Fig. 1c–d).

### 2.2 Refined classification of malignant cells using graph-based smoothing

A key step in identifying target combinations that selectively eliminate leukaemic cells while sparing healthy haematopoietic stem cells (HSCs) was the reliable distinction between malignant and nonmalignant cells within each pAML sample. To this end, we adapted the previously described occupancy score, a measure of the fraction of a patient’s cells assigned to a given cluster, as a proxy for malignancy[17, 28]. This approach is consistent with the observation that cells from the same patient tend to cluster together (Fig. 1c–d) separately from cells from other patients or healthy donors. One limitation of the occupancy score is its dependence on an arbitrary threshold to distinguish healthy and malignant cells, which is inherently difficult to define a priori.

To address the limitations of the occupancy score, we merged and clustered cells from one AML sample together with healthy controls (two adult bone marrow and one paediatric bone marrow samples). Cord blood was excluded due to differences between its HSPCs and those from paediatric or adult bone marrow. To further improve classification, we developed a graph-based approach independent of clustering resolution. For each cell, we assessed its k-nearest neighbors (KNN): if it consistently connected to healthy cells across multiple rounds of label propagation (KNN smoothing), we classified it as healthy. Cells that did not connect to any healthy population after all iterations were classified as malignant (Fig. 1e). In some patients (e.g. AML10) certain healthy progenitors or immune cells also exhibited high occupancy despite clustering with healthy counterparts, likely reflecting AML-induced transcriptional changes (Fig. 1e), highlighting a key limitation of relying solely on the occupancy score for malignant cell classification.

Although fusion transcripts are challenging to detect in scRNA-seq due to dropout and short-read lengths, we identified high-confidence patient-specific fusions (Extended Data Fig. 3 and Supplementary Table 4) that were restricted to cells predicted to be malignant (Fig. 1f). To further evaluate our approach, we benchmarked it against occupancy-based classification across all samples combined (Leiden resolution = 2), which resulted in a higher rate of false negatives—malignant cells harbouring fusions but annotated as normal—highlighting the improved sensitivity of our method (Fig. 1g). Malignant cells defined by our classifier also displayed significantly higher “blast score” than healthy cells from patients, based on a previously published gene signature for paediatric AML[16] (Fig. 1h).

We further applied inferCNV to validate our malignant cell classification in patients with large copy number alterations. In all cases, the inferred copy number profiles recapitulated the expected cytogenetic abnormalities, even revealing additional copy number changes consistent with clonal evolution in one patient at relapse (Extended Data Fig. 4). Using our malignant cell classification to compare paired longitudinal samples revealed relapse-associated transcriptional remodelling, including reduced myeloid differentiation, broadly consistent with previous reports[9], although the limited number of longitudinal samples precluded definitive conclusions (Extended Data Fig. 5) [16].

Collectively, the strong concordance between predicted malignant cells, patient-specific fusion events, inferCNV profiles, blast scores, and paired longitudinal clustering supports the robustness of our malignant cell classification strategy.

### 2.3 Healthy reference-based mapping reveals pAML lineage-biased states linked to cytogenetics

To explore how immunophenotypic heterogeneity within the malignant compartment affects the repertoire of potential therapeutic targets, particularly for immature, LSC-like populations, we mapped AML cells onto a healthy haematopoietic reference built from our own control samples (Extended Data Fig. 6). This allowed us to annotate malignant cells according to their closest normal counterpart and to stratify patients into three groups based on their dominant immunophenotype: primitive-like, GMP-like, and erythroid/MK-like (Fig. 2a–b). These transcriptional identities showed broad patterns of association with cytogenetic subtypes, consistent with prior studies: KMT2A rearrangements were enriched in GMP-like states, RUNX1::RUNX1T1 and CBFB::MYH11 in primitive-like states, and GLIS2::CBFA2T3 cases in erythroid-like states (Fig. 2c). Immunophenotypic assignments were further supported by CITE-seq surface protein expression profiles (Fig. 2d).

**Fig. 2.**
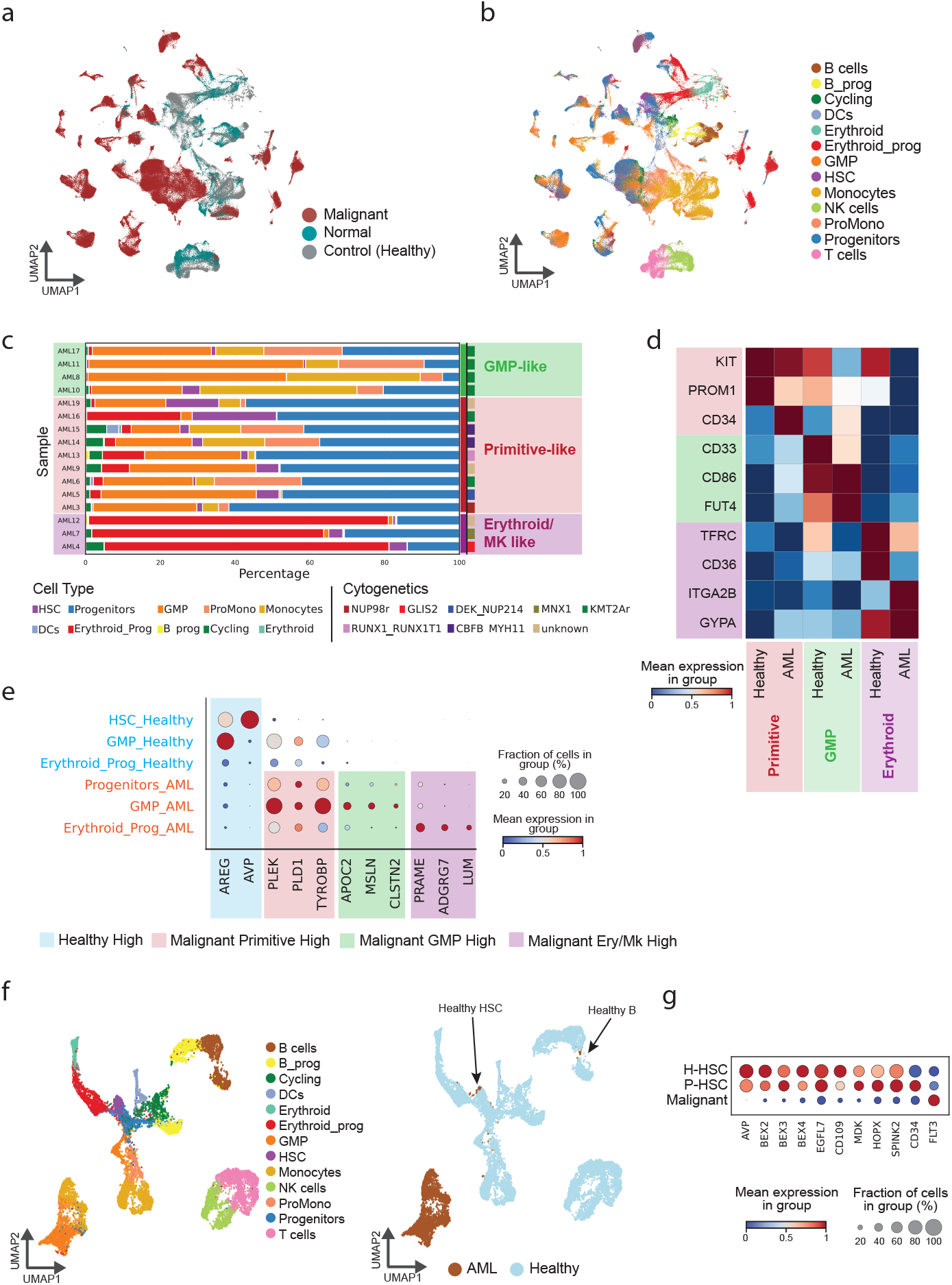
Healthy reference-based mapping reveals pAML lineage-biased states. **a**. UMAP of all single cells in our cohort, coloured by malignant cell and normal cell classification (pAML patients) and sample of origin (healthy controls 2 BM, 2 CB, 1 PBM). **b**. UMAP showing cell type labels across healthy and pAML samples in the cohort. **c**. Bar graph showing cell type contributions within each sample and cytogenetic group. Samples are grouped and labelled by their most enriched cell type as Granulocyte and Monocyte Progenitor (GMP)-like; Primitive-like (enriched for cells labelled as stem or multipotent progenitor cells); erythroid progenitors, and megakaryocyte progenitor (MK)-like cells. **d**. Heatmap showing surface protein levels (antibody-derived tag, ADT) for immunophenotypic markers distinctive of each of the three immunophenotypic groups, across healthy and AML samples. **e**. Dot plot showing RNA expression of selected markers in AML vs healthy samples within the three immunophenotypic groups. The panel includes genes encoding surface proteins, as well as markers such as AREG and AVP, informative for identifying healthy HSCs used in **f–g. f**. UMAP clustering of one AML patient (AML8) and healthy controls, highlighting the identified healthy HSC (AVP^+^) cluster among AML cells. **g**. Dot plot comparing the expression of HSC marker genes between patient-derived HSCs (P-HSCs; patients with *>*30 HSC cells) and healthy HSCs (H-HSCs), relative to malignant cells.

However, given the limited number of samples within individual cytogenetic sub-groups, these observations should be interpreted as trends rather than definitive subtype-specific associations.

Comparison of malignant populations with their healthy counterparts revealed immunophenotype-specific markers. Specifically, APOC2 and MSLN were enriched in GMP-like AML cells, TYROBP and PLEK in GMP and primitive-like AML cells, and PRAME and ADGRG7 in erythroid-like AML cells. Notably, AVP expression was restricted to healthy HSCs, while AREG showed GMP-specific e xpression in healthy bone marrow (Fig. 2e). Through per-patient clustering and KNN smoothing, we identified a rare subset of HSCs from AML samples that clustered with healthy HSCs and lacked detectable fusion events. Interestingly, these cells expressed markers typically restricted to healthy bone marrow HSCs (Fig. 2f). Among these were HOPX, a gene implicated in HSC stemness and quiescence, and EGFL7, which supports HSC survival. Furthermore, CD109 showed low expression in malignant cells but high expression in both healthy HSCs and the patient-derived HSC subset, suggesting potential utility for future strategies aimed at sparing healthy HSCs, for example, as a negative marker in logic-gated CAR designs (Fig. 2g).

While reference-based immunophenotypic labelling provides a useful framework to position malignant cells within the haematopoietic hierarchy, this approach may limit the identification of immature, LSC-like populations that are functionally relevant but deviate from healthy immunophenotypes. Because these labels rely on a healthy reference, they can obscure transcriptional programs shared across cytogenetic backgrounds yet masked by immunophenotypic divergence.

### 2.4 Unbiased identification of LSC-like transcriptional programs conserved across paediatric AML patients

Because healthy-reference anchoring can obscure shared stem-like programs, we shifted to an unbiased strategy aimed at resolving the full spectrum of malignant cell states present in pAML samples (Fig. 3a). We applied non-negative matrix factorization (NMF) to extract gene expression key nodes from the malignant compartment of each patient. This method allows the identification of co-regulated genes, or transcriptional programs within heterogeneous cell populations, without relying on reference datasets, and allows the unbiased characterisation of cell states independent of proximity to healthy cell types. We set the number of programs in NMF to six from each sample and scored all programs against a curated collection of published LSC and blast gene expression signatures[10–13, 16, 18] (Methods). Programs with consistently low weights across cells (*<*0.1), interestingly, also showed no correlation with either LSC or blast signatures and were removed from the analysis. In contrast, programs that positively correlated with LSC score, consistently anticorrelated with the blast score (Fig. 3b). Clustering all programs using the top 50 genes from each patient-specific program revealed two conserved functional modules: one enriched for stem-like features and strongly associated with LSC scores (mean Jaccard similarity across patients = 0.15), and another, more heterogeneous module associated with blast scores (mean Jaccard similarity across patients = 0.09), likely reflecting underlying cytogenetic variation (Fig. 3c). Since clustering based on highly variable genes produced patient-specific clusters that do not capture the hierarchical organisation of AML (Fig. 1d), we decided to restrict the analysis to genes from the LSC–blast axis and perform k-means clustering to unravel conserved gene programs shared across patients.

**Fig. 3.**
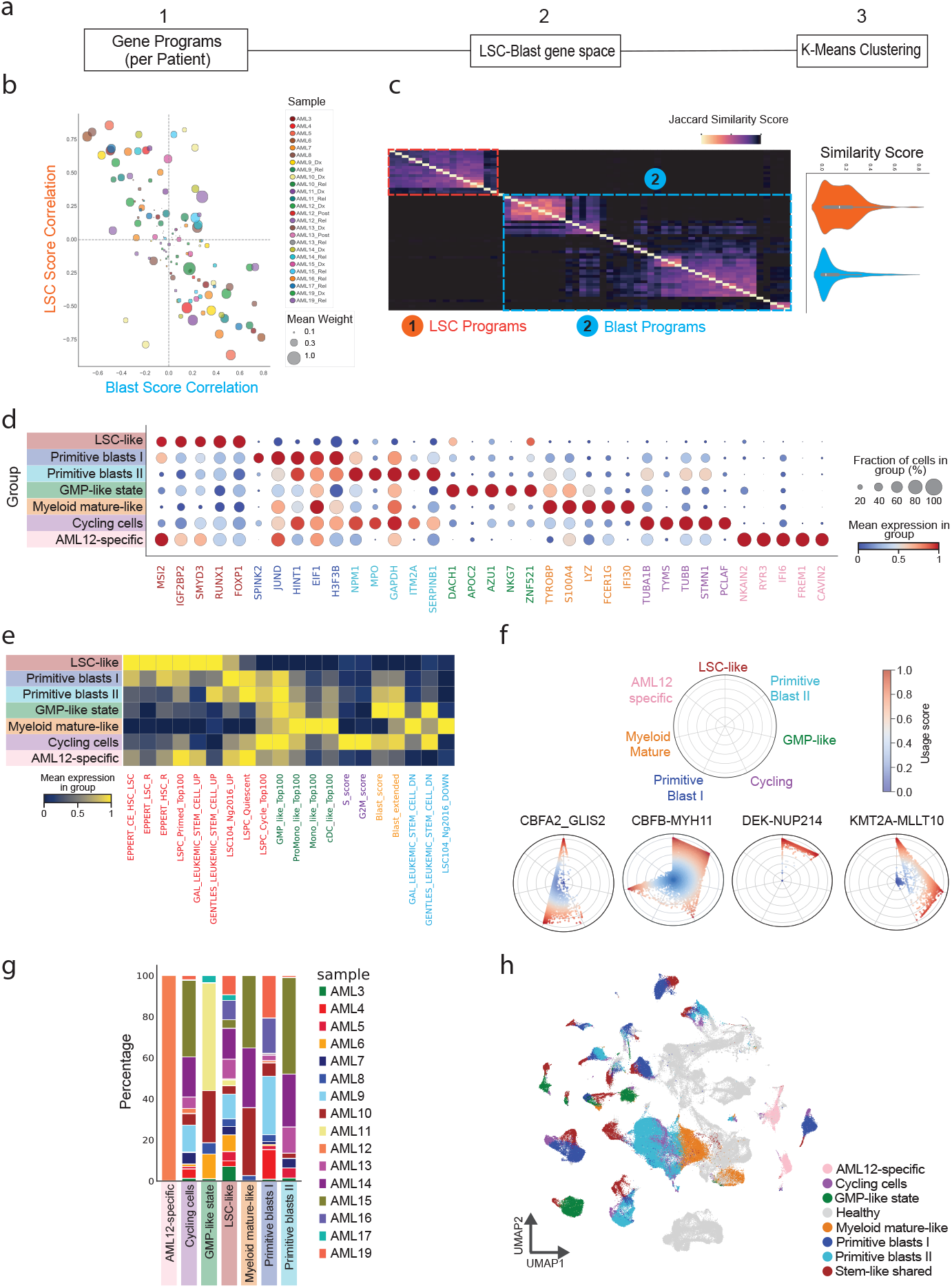
cNMF uncovers conserved stem-like programs across pAML. **a**. LSC identification workflow: per-patient gene programs are identified, programs with low LSC–blast correlation are filtered, and remaining program genes are aggregated to cluster all malignant cells into LSC populations. **b**. NMF-derived transcriptional programs from individual patients plotted by LSC and blast scores (derived from published signatures; see Methods). Points are coloured by sample of origin; size indicates mean program weight across cells. **c**. (left) Jaccard similarity heatmap of programs from panel **a** with weight *>*0.1, clustered by top 50 genes per program, revealing two groups corresponding to LSC and blast programs. (right) Violin plot of Jaccard similarity scores for LSC (orange) and blast (blue) programs. **d**. Dot plot of RNA expression of selected markers across seven conserved cell states identified by *K*-means clustering. Circle size: fraction of expressing cells; colour: mean expression. **e**. AUCell scoring heatmap validating the seven cell states from panel **c**, using published signatures for LSC upregulated genes (red), progenitor/myeloid cells (green), cell cycle (purple), AML (yellow), and LSC downregulated genes (blue). **f**. Radar plots showing enrichment of the seven programs by cytogenetic class: CBFA2T3::GLIS2 (*n* = 1), CBFB::MYH11 (*n* = 2), DEK::NUP214 (*n* = 1), and KMT2A::MLLT10 (*n* = 2). **g**. Sample contribution to each of the seven programs. **h**. UMAP of the full cohort coloured by the seven gene programs.

We identified seven distinct malignant cell states spanning a continuum from LSC-like to GMP-like and monocytic-like cells (Fig. 3d). At the apex was an LSC-like state characterised by high expression of self-renewal–related genes (MSI2, IGF2BP2, RUNX1). Among the blast populations, we annotated Primitive blast I, which expressed stemness-associated genes linked to poor prognosis (e.g., SPINK2, JUND, HINT1), whereas Primitive blast II and GMP-like blasts were enriched for granulocytic progenitor–associated genes (MPO, AZU1), together with metabolic regulators (DACH1, APOC2). Monocytic-like blasts displayed a mature myeloid profile with terminal differentiation markers (S100A4, TYROBP, LYZ, FCER1G). The cycling blast population was defined by high expression of mitotic and cytoskeletal genes (e.g., TUBA1B, TYMS, TUBB, and PCLAF). We also detected a unique patient-specific cluster in AML12, suggestive of transcriptional features not shared with any other patients in our cohort. The annotations of the malignant cell states were validated by AUCell scoring of published LSC, blast, and progenitor gene signatures. The cells classified as LSC-like showed high enrichment for LSC signatures (mean = 0.170 vs. 0.141 for the rest of the malignant population; Mann–Whitney *U* = 2.10 *×* 10^9^, FDR *<* 1 *×* 10^*−*300^) and significantly lower enrichment for blast signatures (mean = 0.109 vs. 0.263; Mann–Whitney *U* = 5.67 *×*10^8^, FDR *<* 1 *×* 10^−300^). In contrast, blast programs, particularly Primitive blast I–II, retained some LSC signature enrichment but also showed increasing enrichment for blast signatures along the differentiation axis (Fig. 3e). The relative contributions of the different blast programs varied depending on cytogenetic subtype, with CBFB::MYH11 fusions enriched in Primitive blast II and mature myeloid states, GLIS2-rearranged cases in Primitive blast I, and KMT2A::MLLT10 in GMP-like states. In contrast, the LSC-like program showed a conserved contribution across almost all patients, irrespective of cytogenetics and immunophenotypes (Fig. 3f–h), facilitating the identification of broadly applicable LSC-directed targets.

### 2.5 Surface antigen landscape of LSC-like cells in pAML

Having identified recurrent malignant cell states across paediatric AML, we next leveraged CITE-seq–derived surface protein data to identify relevant CAR T targets, with a particular focus on the LSC-like compartment. To prioritise surface antigens, we employed expressing cell fraction (ECF) to capture the breadth of expression across cell populations. This approach allowed us to avoid prioritising highly expressed but non-homogeneous targets[29]. From an initial panel of 69 detectable ADTs, we short-listed 19 candidates exhibiting ECF *>* 0.2 in LSC-like cells while maintaining ECF *<* 0.5 in healthy haematopoietic stem cells (HSCs). Examples of best performing targets included CLEC12A (ECF_LSC-like_ = 0.85, ECF_HSC_ = 0.3), CD84 (ECF_LSC-like_ = 0.44, ECF_HSC_ = 0.01), and IL1RAP (ECF_LSC-like_ = 0.72, ECF_HSC_ = 0.01). Conversely, targets such as CD244, CD99, and ITGAL demonstrated high coverage in both LSC-like and HSC populations (ECFs *>* 0.8) and were excluded due to potential toxicity (Fig. 4a and Supplementary Table 5). Our analysis also confirmed previously described HSC-depleting targets, including CD33 and CD123 (IL3RA), as well as “group-specific” candidates such as CD70 and MSLN, which displayed variable ECF in LSC-like cells[22], likely reflecting preferential expression in specific cytogenetic subtypes. Notably, the top-performing targets identified in LSC-like cells also showed high ECF in the blast population (ECF *>* 0.5), indicating that targeting these antigens could simultaneously eliminate both LSC-like cells and blasts(Fig. 4b). Furthermore, comparison of RNA-derived ECF with ADT-derived ECF highlighted the necessity of the ADT modality for target discovery, as the RNA modality was sparse and failed to exceed 0.4 ECF across all groups. One exception was ADGRE2, which showed poor concordance between RNA and protein levels, likely reflecting a marker-specific limitation of the CD312 ADT . However, none of the identified targets alone achieved complete coverage of all LSC-like cells in our cohort (*>*0.85), motivating the exploration of combinatorial targeting strategies to maximise therapeutic reach.

**Fig. 4.**
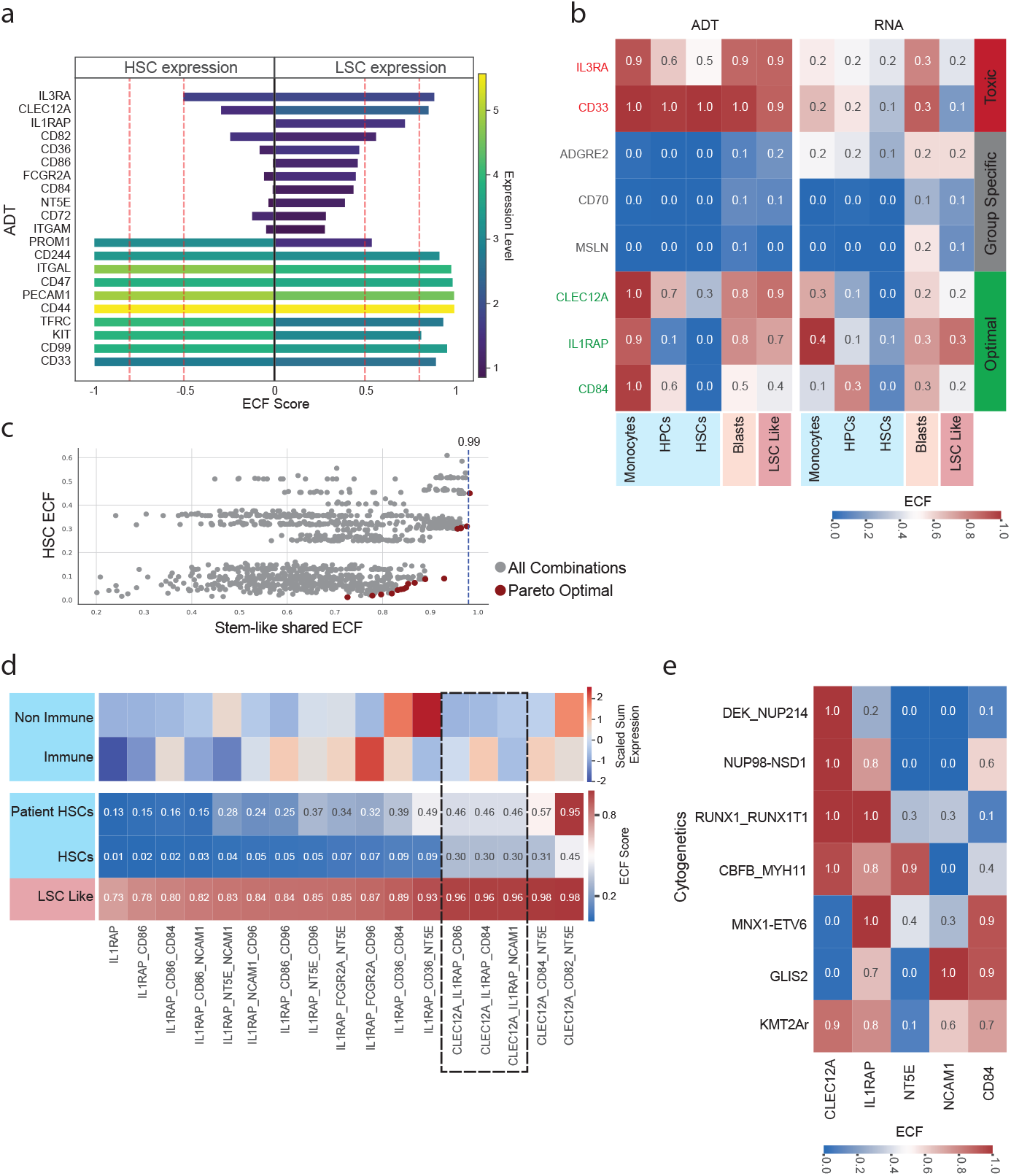
Surface antigen landscape of LSC-like cells in pAML. **a**. Top surface markers ranked by antibody-derived tag (ADT) expressing cell fraction (ECF) in healthy donor HSCs (left) versus LSC-like cells from all pAML patients (right). The 10 markers with the highest ECF in LSCs are displayed at the top, while markers most highly expressed in HSCs are shown at the bottom. Bar lengths indicate ECF, and colour reflects ADT expression levels. **b**. Heatmap showing ADT- and RNA-derived ECF for representative surface markers classified as “toxic” (high ECF in HSCs), “group-specific” (restricted to a subset of patients), or “optimal” (high ECF across most patients), displayed across LSC-like cells, blasts, and healthy compartments (monocytes, progenitors (HPCs), and HSCs). **c**. Scatter plot showing single and combinatorial (2–3) non-HSC-toxic ADT markers. Axes represent LSC coverage (LSC ECF) versus HSC coverage (HSC ECF). Pareto-optimal combinations that maximise LSC targeting while minimising HSC coverage are highlighted in red. **d**. Heatmap of all Pareto-optimal target combinations, showing collective combination ECF across healthy HSCs (from healthy donors and patient-derived HSCs in our cohort) as well as summed expression in immune and non-immune tissues from the Tabula Sapiens dataset. **e**. Heatmap showing ECF values for all optimal markers included in the three top-performing combinations (panel **d**), across individual patients in our cohort, stratified by cytogenetic subgroup.

We applied multi-objective optimisation to identify combinations of up to three targets from our optimal target list that maximised LSC coverage while minimising HSC toxicity (Fig. 4c and Supplementary Table 6). This approach yielded 17 Pareto-optimal combinations, which we systematically evaluated against healthy HSCs from both our in-house healthy haematopoietic reference and patient cohort, and a compendium of healthy immune and non-immune tissues from the Tabula Sapiens single-cell RNA-seq atlas. Combinations containing CLEC12A exhibited particularly broad coverage of the LSC-like cells, with only a modest increase in HSC targeting in both healthy donor and patient-derived cells (Fig. 4d, highlighted). For example, combinations such as NT5E CD82 CLEC12A and NT5E CD84 CLEC12A reached near complete coverage of the LSC-like cells (ECF = 0.98) but also showed high ECF in patient-derived HSCs, suggesting potential relevance for myeloablation before allogeneic haematopoietic stem cell transplantation rather than stand-alone therapies. When accounting for HSCs and normal tissue toxicity, the most effective combinations were CD86, CD84 or NCAM1 together with IL1RAP and CLEC12A (ECF = 0.96) (Fig. 4d). To investigate whether the predicted efficacy was affected by pAM L cytogenetic class or influenced by the composition of our cohort, we next mapped the expression of the Pareto-selected targets across cytogenetic groups. CLEC12A demonstrated near-complete coverage in patients with DEK::NUP214, NSD1::NUP98, RUNX1::RUNX1T1, and CBFB::MYH11 fusions. By contrast, NT5E (CD73) exhibited cytogenetic class-specific expression, almost exclusively in C BFB::MYH11 patients, while NCAM1 (CD56) expression was largely restricted to GLIS2-rearranged patients. IL1RAP showed broad coverage across all cytogenetic groups, with high prevalence in RUNX1::RUNX1T1 patients and those with MNX1 fusions. CD84 was also highly expressed in both GLIS2- and MNX1-rearranged groups. Notably, no single target achieved full coverage of KMT2Ar patients, highlighting further heterogeneity in the LSC-like cells even within the same cytogenetic group (Fig. 4e). To resolve heterogeneity in highly variable groups such as the KMT2Ar and to define minimal sets of targets required to achieve effective L SC-like c overage, we examined the diversity within the LSC-like compartment.

### 2.6 Distinct LSC-like compartments underpin target heterogeneity

While the antigen prioritisation above was performed at the protein level, understanding why no single marker achieved universal coverage required us to first resolve the underlying transcriptional heterogeneity of LSC-like cells. To this end, we clustered LSC-like cells from all patients based on RNA modality, and we observed that despite a small fraction of cells clustered in a patient-specific manner, the majority formed shared clusters across patients (Fig. 5a–b). Within the shared LSC-like core transcriptional program, we identified three main subgroups, likely reflecting underlying immunophenotypic diversity. Primitive LSC-like cells (P-LSCs) expressed canonical stem/progenitor surface markers including CD34, CD117, and CD133. Monocytic-like LSCs (M-LSCs) lacked CD34 expression but expressed CD4, CD11a, and CD36, consistent with prior reports[19, 30]. A third group, termed erythroid-like LSCs (E-LSCs), was characterised by CD41 and CD71 expression, indicative of erythroid/megakaryocytic lineage.

**Fig. 5.**
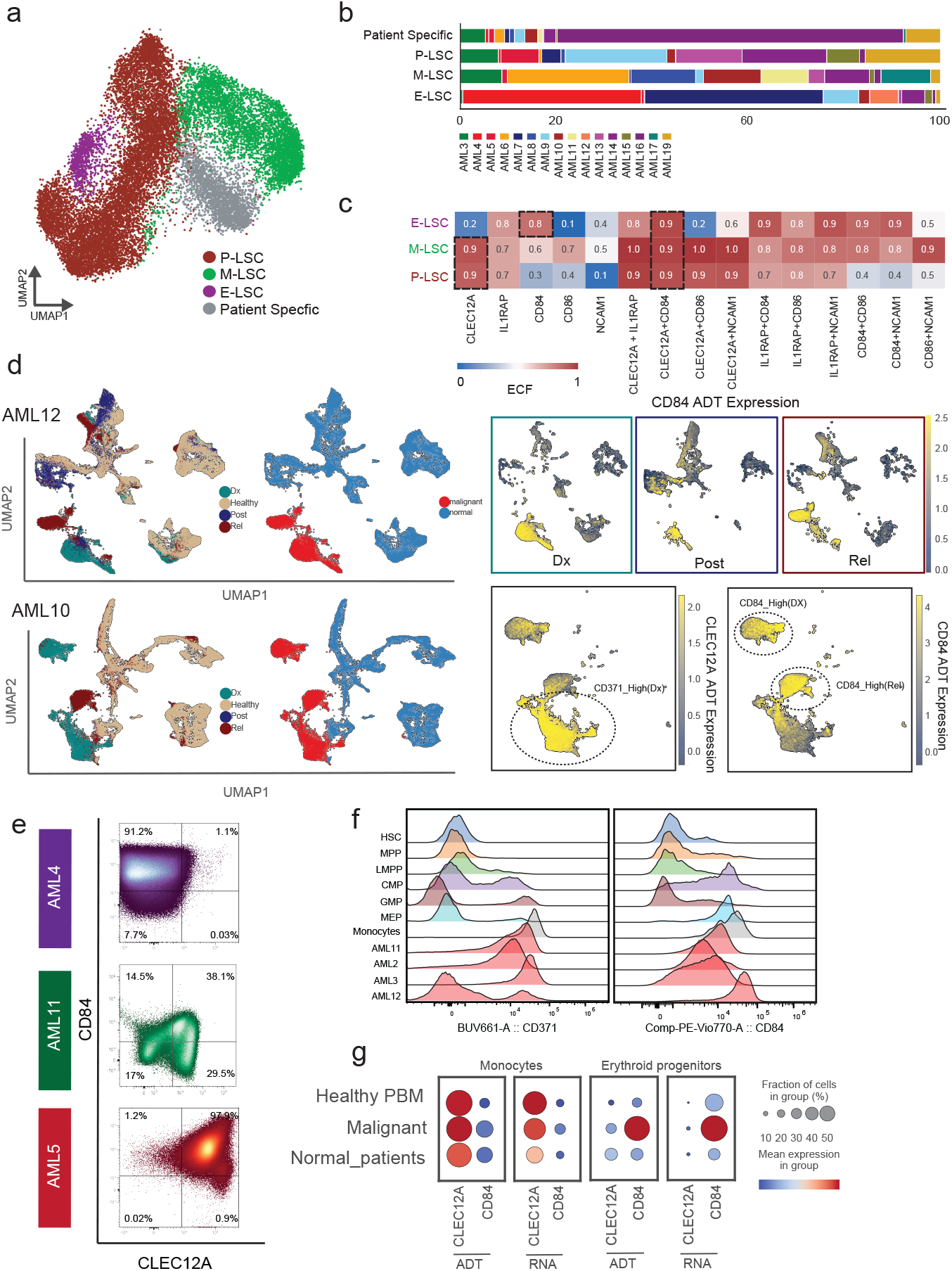
Distinct LSC-like compartments underpin target heterogeneity. **a**. UMAP of LSC-like cells from all patients, revealing three subtypes: primitive (P-LSC), monocytic (M-LSC), and erythroid-like (E-LSC), with a small fraction of patient-specific clusters. **b**. Sample contribution to each LSC subtype. **c**. Expressing cell fraction (ECF) for single markers from the optimal Pareto set (Fig. 4d) and their combinations across LSC subtypes. **d**. UMAPs of AML12 (top) and AML10 (bottom) coloured by timepoint (Dx: diagnosis, Post: remission, Rel: relapse), malignant/normal identity (KNN smoothing), and CD84 ADT expression per timepoint, showing consistent CD84 enrichment in malignant cells across disease course. In AML10, a CD371-high population is highlighted at Dx, and two CD84-high populations at Dx and relapse, marking antigenically distinct subpopulations across disease stages. **e**. Flow cytometry of CD45-positive bone marrow mononuclear cells from three patients representing each LSC subtype: AML4 (GLIS2-rearranged, E-LSC), AML11 (KMT2A::MLLT10, M-LSC), and AML5 (DEK::NUP214, P-LSC). **f**. Ridge plots of CD371 (left) and CD84 (right) expression by flow cytometry across haematopoietic populations (HSC, MPP, LMPP, CMP, GMP, MEP, monocytes) in four healthy bone marrow donors (top) and four paediatric AML patients—AML11, AML2, AML3, AML12 (bottom). **g**. Dot plots of ADT and RNA expression of CLEC12A and CD84 in monocytes and erythroid progenitors across healthy paediatric bone marrow (Healthy PBM), malignant, and normal bystander cells (Normal_patients) from paediatric AML patients. Dot size: fraction of expressing cells; colour: mean expression (red: high, blue: low).

To place these pAML LSC subgroups within the broader adult LSC landscape, we compared their transcriptional profiles with published adult AML LSC signatures (Extended Data Fig. 7f). P-LSCs showed the highest similarity to previously described adult LSC states, particularly the LSC-primed and quiescent subpopulations described by Zeng and colleagues[18]. In contrast, M-LSCs were enriched for adult GMP-like, promonocytic, and monocytic stem-cell programs, including the monocytic LSC (m-LSC) state described by Sango et al.[20] and displayed low BCL2 and high MCL1 expression, consistent with transcriptional features associated with venetoclax resistance in adult AML (Extended Data Fig. 7c)[19, 20]. Although adult-derived LSC scores were generally higher in P-LSCs, E-LSCs also retained enrichment for a subset of primitive and stemness-associated signatures, indicating that stemness-associated programs are not restricted to the P-LSC compartment (Extended Data Fig. 7f). Together, these findings indicate substantial overlap with adult AML while highlighting distinct LSC states in paediatric disease.

We then assessed the expression of the Pareto-optimal targets across these three LSC-like subtypes, considering both single markers and combinations (Fig. 5c). As single markers, CLEC12A alone showed ∼0.9 ECF in both P-LSCs and M-LSCs, but only 0.2 in E-LSCs, suggesting a risk of antigen-negative escape in patients enriched for this LSC subtype. In contrast, CD84 emerged as the best-performing target for E-LSCs (ECF *>* 0.8). Among combinations, CLEC12A_IL1RAP provided the highest coverage of P-LSCs, whereas CLEC12A_CD86 was optimal for M-LSCs. Ultimately, the CLEC12A/CD84 combination achieved the most balanced coverage across P-, M-, and E-LSCs.

We then annotated each patient by their predominant LSC-like subtype; although multiple subtypes often coexisted, one typically predominated. Notably, the same antigen co-expression pattern was evident across all malignant cells (LSC-like and blasts), suggesting that the same target combinations would likely be effective against the broader malignant compartment (Extended Data Fig. 7g). Notably, in a patient with predominant E-LSC identity (AML12), CD84-positive malignant cells were detectable at diagnosis and CD84 expression remained stable across all longitudinal samples. In addition, the utility of CD84 as a target extends beyond the E-LSC context. In a second patient with KMT2Ar AML (AML10), a subset of CD84-high malignant cells present at diagnosis persisted at relapse, further supporting the rationale for combinatorial targeting across multiple LSC states (Fig. 5d).

We next evaluated the expression of these three targets across healthy tissues. Analysis of publicly available bulk and single-cell RNA-seq datasets (Human Protein Atlas and Tabula Sapiens[31, 32]), together with our healthy CITE-seq reference, revealed highly concordant expression patterns (Extended Data Fig. 8). CLEC12A was largely restricted to the myeloid differentiation trajectory from CMP/GMP to mature monocytes, IL1RAP was predominantly expressed on mature monocytes, and CD84 was enriched in CMP/MEP populations and monocytes, whereas all three markers showed minimal expression in HSCs and multipotent progenitors. While CLEC12A and CD84 expression remained largely confined to immune compartments, IL1RAP displayed elevated expression in hepatocytes, raising potential concerns regarding liver toxicity and narrowing subsequent analyses to CLEC12A and CD84 (Extended Data Fig. 8).

Flow cytometry of healthy bone marrow (n = 4) confirmed the lineage-specific expression patterns of CLEC12A and CD84 observed by CITE-seq. Moreover, quantitative antigen-density measurements strongly correlated with CITE-seq ADT expression across healthy haematopoietic populations (Extended Data Fig. 8f), confirming monocytes and MEPs as the healthy compartments with the highest physiological expression of CLEC12A and CD84, respectively, and supporting the quantitative value of the CITE-seq protein measurements. We next validated these findings in six paediatric AML samples spanning the major LSC subtypes identified in our cohort. Consistent with the single-cell analyses, E-LSC-enriched samples showed low or absent CLEC12A expression with preferential CD84 coverage, whereas P/M-LSC–enriched samples expressed both markers, with CLEC12A providing broad coverage and CD84 contributing complementary coverage (Fig. 5e-f and Extended Data Fig. 8e). Importantly, antigen density in AML reflected the subtype-specific CITE-seq findings, with high CLEC12A expression in P/M-LSC–enriched AML and high CD84 expression in the E-LSC–enriched AML sample (Extended Data Fig. 8f).

Using CITE-seq, we compared RNA and ADT expression of malignant cells with healthy cells from both healthy donors and residual normal cells within AML samples. Residual healthy populations showed comparable or lower CLEC12A and CD84 RNA and ADT expression than the corresponding healthy donor populations, suggesting that the leukaemic bone marrow microenvironment does not increase target expression in non-malignant cells (Fig. 5g). Together, these findings support the robustness of the CLEC12A/CD84 combination across heterogeneous paediatric AML cell states.

### 2.7 Protein prediction and bulk transcriptomics confirm LSC-type stratification and link it to target landscapes

To validate the robustness of our LSC-like program and test whether the target profiles identified in our cohort could be validated in independent datasets, we analysed the non-CD34-enriched scRNA-seq dataset of paediatric and adult AML developed by Lasry et al.[28]. We employed a healthy–malignant classifier trained on our cohort to identify malignant cells in the external samples, followed by clustering based on NMF-derived gene programs obtained from our cohort. In line with our dataset, we consistently identified an LSC-like population with the highest LSC scores across the majority of samples (Extended Data Fig. 9). As this external cohort included FAB and flow cytometry data for some patients, we selected representative cases with available FAB classification and/or clinical immunophenotype and mapped the defined LSC-like cells onto our LSC reference (Fig. 6a and Extended Data Fig. 10f). The mapping revealed distinct LSC subtypes. Notably, LSCs from three KMT2Ar cases exhibited clear subtype-specific patterns: AML052 (KMT2A::MLLT10, FAB M7) mapped exclusively to the E-LSC cluster, AML048 (KMT2A::MLLT3, CD34^−^ myeloid phenotype) mapped to the M-LSC cluster, and AML025 (KMT2A–PTD, CD34^+^ progenitor phenotype) mapped to the P-LSC cluster. In each case, the LSC-type assignment was consistent with FAB classification and/or clinical immunophenotype, confirming the ability of our framework to capture distinct LSC subtypes.

**Fig. 6.**
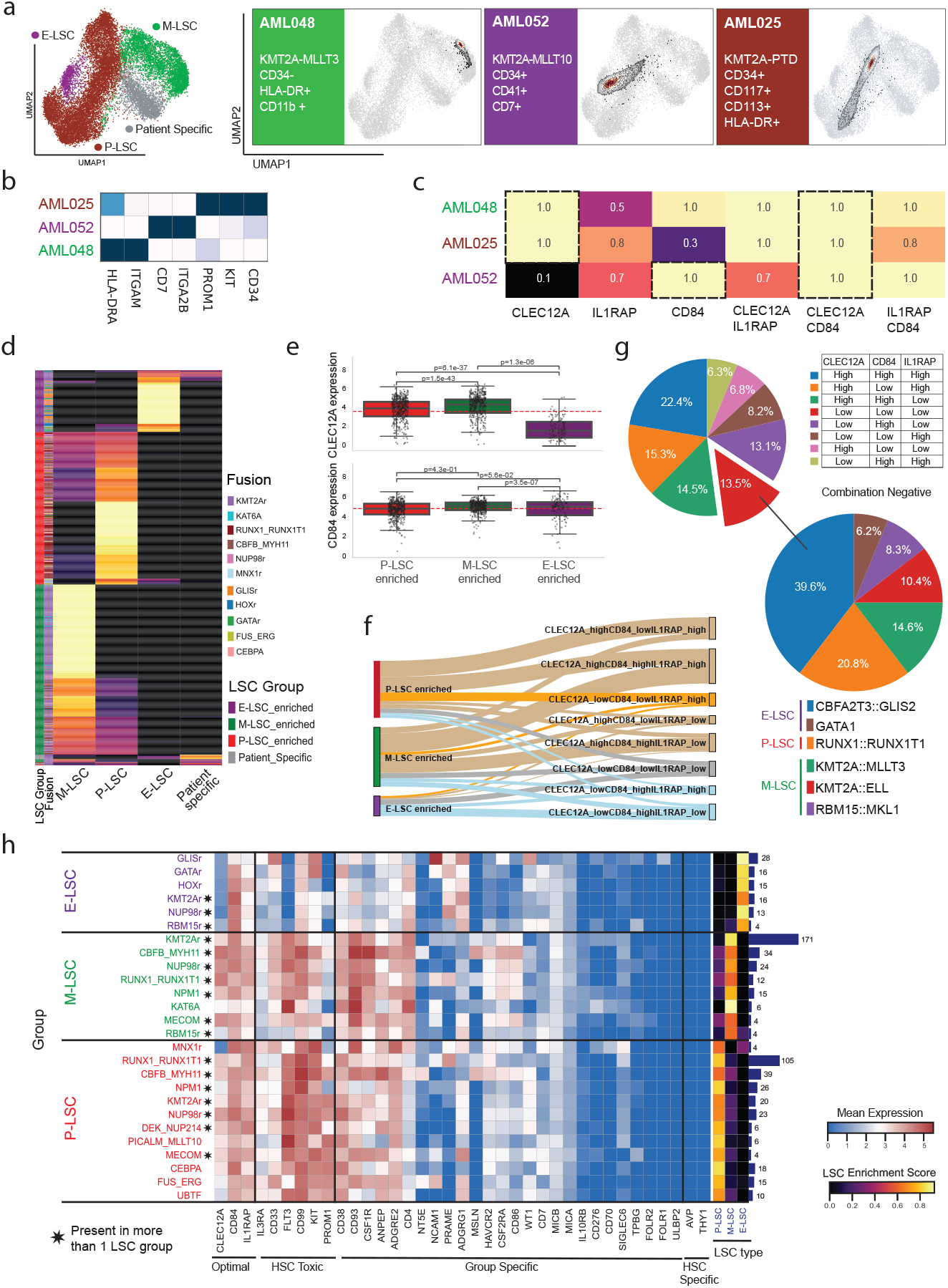
Protein prediction and bulk RNA validate LSC-type targets. **a**. LSC reference UMAP (P-LSC, M-LSC, E-LSC) with projected LSC-like cells from three KMT2A patients in the Lasry dataset, alongside their clinical immunophenotypes as reported in Lasry et al. **b**. Predicted ADT expression for the same patients via scTranslator, recapitulating reported clinical immunophenotypes. **c**. Predicted ADT-derived ECF for CLEC12A, CD84, and IL1RAP across the three KMT2A patients. **d**. Heatmap of the St. Jude bulk RNA cohort (*n* = 988) showing patient-level deconvolution by P-, M-, and E-LSC gene signatures. Left bars indicate dominant LSC type and cytogenetic/molecular classification; colour scale represents LSC-type signature expression. **e**. CLEC12A and CD84 RNA expression across the St. Jude cohort stratified by dominant LSC type. Kruskal–Wallis test: CD84 *p* = 3.32 *×* 10^−6^, CLEC12A *p* = 8.55 *×* 10^−48^. Pairwise Mann–Whitney U-tests: CD84 M-LSC vs P-LSC *p* = 3.46 *×*10^−7^; CLEC12A E-LSC vs M-LSC *p* = 1.51 *×*10^−43^, E-LSC vs P-LSC *p* = 6.05 *×*10^−37^, M-LSC vs P-LSC *p* = 1.31 *×*10^−6^. **f**. Sankey plot of high vs low RNA expressors for CLEC12A, IL1RAP, and CD84 by dominant LSC type in the St. Jude cohort. **g**. Pie charts showing the proportion of marker-positive and marker-negative patients (RNA) across the full cohort (upper left), with cytogenetic subgroup and LSC-type enrichment among triple-negative patients (lower right). **h**. Heatmap of St. Jude patients by cytogenetic/molecular classification (Umeda et al.), subdivided by dominant LSC type, showing RNA expression of optimal targets (CLEC12A, CD84, IL1RAP), HSC-toxic markers, and additional CAR T candidate antigens. Right bar plot shows LSC-type enrichment scores per patient group.

Since the dataset by Lasry et al. lacked several key ADTs in the CITE-seq panel, we employed scTranslator[33], a pre-trained generative model, which we fine-tuned on our cohort to predict the missing ADTs in the LSC-like cells of the external dataset. The model accurately predicted canonical immune microenvironment markers, supporting its ability to generate reliable predictions (Extended Data Fig. 10). We next applied it to the same three KMT2Ar patients described above and found that predicted proteins well recapitulated the immunophenotypic annotations from clinical data (Fig. 6b). Predicted abundance of our key targets confirmed complementary profiles, with C LEC12A s howing high c overage in P - and M -LSCs, but absent in E-LSC-enriched case, where CD84 provided compensatory coverage. The combination of CLEC12A and CD84 achieved the highest and most balanced coverage in all three patients, further reinforcing our findings from the internal cohort in an independent dataset (Fig. 6c).

Given the limited number of patients and cytogenetic subgroups available in our own and other scRNA-seq cohorts, we extended our LSC typing framework to an external bulk RNA-seq from the St. Jude cohort. We applied LASSO regression for deconvolution using the three identified L SC signatures ( Supplementary Table 7) and stratified patients into three main groups: an E -LSC group enriched mostly for GLIS2r, HOXr, GATAr, subsets of KMT2Ar, and NUP98r patients; an M-LSC group enriched for KMT2Ar cases with subsets of NPM1 and NUP98r patients; and a P-LSC group enriched for RUNX1r, CBFB::MYH11, NUP98r, DEK::NUP214, CEBPA, FUSr, and subsets of NPM1 and KMT2Ar patients. The E-LSC group was associated with M6–M7 FAB classification, consistent with prior annotations[7]. While most patients showed predominant enrichment for a single LSC subtype, co-occurrence of LSC programs was also observed. In particular, P-LSC and M-LSC signatures frequently co-occurred across multiple cytogenetic groups. By contrast, co-enrichment of P-LSC and E-LSC signatures was largely restricted to MNX1-rearranged cases, consistent with our previous observation that this subgroup exhibits mixed primitive and erythroid-like features (Fig. 6d).

Target mapping across these groups recapitulated single-cell findings, with CLEC12A highly expressed in both P- and M-LSC groups but low in E-LSC group (Kruskal–Wallis test, *p* = 2.69 *×* 10^−34^) where CD84 compensated (Fig. 6e). To further investigate combinatorial targeting, we binarized samples into high (*>*mean) and low (*<*mean) expression for each of the three targets and assessed expression patterns across LSC groups. Notably, CD84 was not only required to compensate for the lack of CLEC12A in E-LSCs but was also essential in certain cases within the M-LSC group. Similarly, IL1RAP, which exhibited the highest ECF in P-LSC–enriched cases, was further shown as a critical co-target to improving patient coverage in this group. This is particularly relevant for effectively targeting C LEC12A-negative P-LSC–enriched cases, which include those associated with RUNX1 rearrangements (Fig. 6f). Around 13% of the pAML St. Jude cohort showed low expression of all three targets, largely originating from GLISr and KMT2Ar patients, mostly corresponding to the E- and M-LSC subtypes. Although we demonstrated by both flow cytometry and our ADT modality that the GLIS2r patient in our cohort is positive for CD84, substantial heterogeneity remains, highlighting the need to identify more uniformly expressed targets across this patient group (Fig. 6g). We next asked whether our LSC typing could resolve heterogeneity within genetically defined p AML classes by subdividing cytogenetic classes according to their dominant LSC type. For instance, KMT2Ar and NUP98r groups both stratified into all three E -, P -, and M -LSC c lasses. In contrast, CBFB::MYH11 and RUNX1r patients mainly split into P- and M-LSC groups, while homogeneous groups such as GLISr patients were only found within the E-LSC class (Supplementary Table 8). We then checked these newly defined groups (cytogenetics + dominant LSC type) against a panel of known AML CAR targets. This integrated classification unravelled target expression patterns otherwise undetectable by cytogenetics alone. While P-LSCs appeared broadly targetable not only with our preferred combination, but also with additional markers such as FLT3 and CD99, and M-LSCs offered additional opportunities, including CSF1R, CD93, and CD4, the E-LSC group emerged as the most challenging to target. While these cells showed consistently low expression of canonical AML targets such as CLEC12A, CD33, IL3RA, and FLT3, potential alternative targets included CD84 and NCAM1 (in the GLISr group), with PRAME and ADGRG1 being selectively upregulated in this subset (Fig. 6h).

Finally, we applied the same approach to the TARGET cohort, which yielded LSC typing and target expression patterns consistent with those observed in the St. Jude (Extended Data Fig. 11 and Supplementary Table 9). Importantly, the availability of survival data in the TARGET cohort allowed us to explore whether LSC classification identified biologically distinct patient groups within established cytogenetic categories. Across the majority of the cytogenetic groups (RUNX1r, NPM1, FLT3-ITD, and CBFB::MYH11 patients), M-LSC enrichment was associated with a trend toward worse survival compared to P-LSC enrichment. Interestingly, the opposite trend was observed in KMT2Ar patients, in which P-LSC enrichment was linked to inferior outcomes (Extended Data Fig. 12). Although these findings were not consistent across all cytogenetic groups and should be interpreted cautiously, they suggest that LSC composition may provide additional biological context within established molecular subtypes.

Together, we showed that across independent scRNA-seq and bulk RNA cohorts (Lasry, St. Jude, and TARGET), our framework consistently recovered LSC-like sub-types, recapitulating their clinical immunophenotypes and linking them to distinct antigen profiles. Notably, even in bulk datasets, despite the lack of LSC enrichment and reliance on RNA modality alone, the complementary role of CD84 in compensating for low CLEC12A expression was confirmed. These findings highlight the robustness of our approach and its potential to provide therapeutic insights beyond cytogenetic classification alone.

## 3 Discussion

Targeting LSCs is essential for durable therapeutic responses in AML. While adult single-cell studies have substantially advanced our understanding of LSC biology, heterogeneity and therapeutic vulnerabilities, paediatric single-cell studies have so far focused primarily on relapse-associated transcriptional remodelling, the tumour microenvironment, or blast-cell differentiation states rather than systematic discovery of LSC-directed immunotherapy targets [9, 16–18]. As a result, a paediatric-specific framework for LSC-directed antigen prioritisation has been lacking.

To overcome this, we systematically evaluated 81 surface antigens by CITE-seq, across our paediatric AML cohort, of which 69 passed quality control and were prioritised through sequential filtering based on LSC enrichment and healthy tissue expression. By integrating surface protein profiling with unbiased transcriptional analysis, we resolved the LSC compartment across immunophenotypically diverse patientspatients, a critical step for defining CAR T targets directed at LSCs. We identified highly conserved gene programs characteristic of LSC-like states, suggesting that, despite the substantial genetic and phenotypic diversity of paediatric AML, stem-like programmes are constrained to a limited number of recurrent biological states, thereby providing the foundation for therapeutic target prioritisation.

LSC-like cells exhibited heterogeneous surface antigen profiles and could be sub-divided into three principal categories: primitive (P-LSC), monocytic (M-LSC), and erythroid-like (E-LSC). The identification of a distinct M-LSC subtype is consistent with recent adult AML studies describing a CD34^−^CD4^+^ monocytic LSC population associated with adverse outcome and venetoclax resistance [19, 20]. In contrast, the E-LSC state has been underrepresented in previous single-cell studies, limiting systematic evaluation of therapeutic targets in erythroid-biased disease [9, 28]. While we did not specifically annotate a lymphoid LSC subtype, CD34^+^ lymphoid progenitor-like states are likely included within the broader P-LSC category. These LSC subtypes frequently coexisted within individual patients in varying proportions. Notably, while P-LSC and M-LSC programmes co-occurred across multiple cytogenetic groups, P-LSC and E-LSC co-occurrence was largely restricted to MNX1-rearranged AML, supporting LSC-state composition as an additional layer of heterogeneity beyond cytogenetics, with potential implications for patient stratification and the design of combinatorial targeting strategies.

While we cannot exclude that our limited cohort composition influenced the LSC types that we identified, expanding our RNA-surface protein-matched datasets may refine this classification further. Nevertheless, these findings suggest that paediatric AML shares core stem-cell programmes with adult disease, while retaining distinct LSC-state distributions and antigen-expression patterns that may have important implications for target prioritization.

Using CITE-seq data from our cohort and models trained to predict protein expression from RNA in external single cell datasets, we mapped surface targets and identified combinations capable of eradicating all LSC states (P-, M-, and E-LSCs). CLEC12Aand CD84 together showed the broadest LSC complementary coverage across LSC compartments, while maintaining favourable healthy-tissue expression profiles. IL1RAP also emerged as a promising LSC-directed target and has demonstrated efficacy in patient-derived xenograft models [34],although its broader expression in hepatocytes may complicate clinical translation (Extended Data Fig. 8d). Target expression varied systematically across LSC states. CLEC12A predominantly covered P- and M-LSCs, whereas CD84 complemented CLEC12A by extending coverage to E-LSCs, where CLEC12A expression was largely absent. The contribution of CD84 extended beyond erythroid-biased disease. CD84-high malignant populations were also observed in P/M-LSC-enriched disease and persisted at relapse, as exemplified by the KMT2A-rearranged AML10 case, supporting combina-torial CLEC12A/CD84 targeting across multiple LSC states.

Importantly, the reproducibility of LSC-state composition and associated target landscapes across independent single-cell and bulk RNA-seq cohorts demonstrates that these conserved stem-like programmes can be identified without paired protein measurements. Integrating LSC-state composition with cytogenetic subtype further refined patient stratification and enabled identification of subtype-specific immunotherapy targets[22], including NCAM1 for GLIS2-rearranged AML, IL1RAP for RUNX1-rearranged AML, and LSC-specific targets such as CD93 for M-LSC and PRAME or ADGRG1 for E-LSC subpopulations.

Functionally, CLEC12A is a C-type lectin receptor with inhibitory signalling activity but no established role in AML cell survival. In contrast, CD84 has recently been shown to support mitochondrial function and antioxidant defence in AML, promoting LSC self-renewal and propagation [35].

CLEC12A and CD84 are both currently being evaluated as CAR targets, including armoured IL-18-secreting CLEC12A-directed constructs[24, 36, 37]. CD84 has recently shown efficacy in xenograft models with relatively low myelotoxicity and has entered early-phase clinical evaluation for relapsed/refractory AML (NCT07471789)[37, 38]. CLEC12A is more clinically advanced, with adult AML trials reporting ∼70%remission rates, albeit with frequent high-grade cytokine release syndrome (CRS) and prolonged cytopenias[24, 39], while paediatric studies indicate remissions with com-paratively lower myelotoxicity[36, 40]. Across paediatric trials, most children achieved short-term remission, though some, particularly NUP98r and erythroid (M6) AML cases showed no remission or early relapse, likely reflecting variable CLEC12A antigen coverage across cytogenetic and immunophenotypic subtypes[36, 40].

Our findings provide a potential biological explanation for these heterogeneous clinical responses. E-LSC-enriched AML exhibits low CLEC12A expression and may therefore be intrinsically less responsive to CLEC12A-directed monotherapy, whereas patients with mixed P- and M-LSC populations are unlikely to be fully covered by a single antigen. Accordingly, combinations not informed by LSC heterogeneity, such as those incorporating CLEC12A with CD33 or CD123[41, 42], are unlikely to fully eradicate E-LSCs while potentially increasing HSC toxicity. By contrast, the complementary expression patterns of CLEC12A and CD84 support their prioritisation as a rational dual-targeting strategy.

In summary, our work establishes the first paediatric-specific framework for LSC antigen prioritisation in AML, demonstrating how LSC-resolved antigen profiling can identify rational target combinations, exemplified b y C LEC12A and C D84, to address LSC heterogeneity in paediatric AML. More broadly, the target prioritisation framework developed here could be easily extended to other haematological and non-haematological malignancies. By combining paired transcriptomic and surface proteomic measurements with predictive modelling this framework can be extended to existing single-cell and bulk RNA-seq cohorts. This approach enables integration of immunophenotype, cytogenetics, and cell state programs into a unified classifier, refining disease subgroups and informing the design of antigen combinations increasingly tailored to individual patient profiles. Ultimately, LSC-informed target selection provides a path toward more precise and effective CAR T strategies, accommodating both inter-patient heterogeneity and the diversity of stem-like compartments within each leukaemia.

## 4 Methods

### 4.1 Antibody–oligonucleotide conjugation for CITE-seq

For non-commercially available targets, we performed in-house antibody– oligonucleotide conjugation. Unconjugated antibodies against Relaxin1 (RXFP1, MAB8898, R&D Systems), Mesothelin (MSLN, 530101, BioLegend), SIGLEC6 (MAB2859, R&D Systems), IL1RAP (MAB676, R&D Systems), CD312/ADGRE2 (W15101A, BioLegend), BCL-2 (BMS1029, Thermo Fisher), KDEL/GPR78 (EPR12668, Abcam), were conjugated with custom oligonucleotides corresponding to feature barcodes of BioLegend TotalSeqC antibodies. Oligonucleotides were ordered from IDT (UK). Conjugations were performed using either the Abcam Oligonucleotide Conjugation Kit (#ab218260) or the 5^*′*^ Feature Barcode Antibody Conjugation Kit – Lightning-Link^®^ (#ab270703), following the manufacturer’s instructions. For most conjugations, 20–100 *µ*g antibody input was used.

### 4.2 Patient samples, antibody staining, and flow cytometric sorting

Paediatric AML samples and their clinical details are listed in Supplementary Table 1. Samples were obtained from the VIVO biobank or Great Ormond Street Hospital (GOSH). Healthy bone marrow (BM) controls were obtained from Lonza. All patients (or their legal guardians) provided written informed consent in accordance with the Declaration of Helsinki for sample collection and research use. BM mononuclear cells (MNCs) were isolated by Ficoll density-gradient centrifugation. Cryopreserved BM MNCs were thawed on the day of the experiment and Fc-blocked using Human TruStain FcX™ (BioLegend, #422301) and Miltenyi FcR-blocking reagent (#130-059-910). Antibody mixes were freshly prepared before each experiment. Following Fc-blocking, cells were incubated with antibody mixes for 30 min on ice, protected from light, washed 1–3 times with PBS containing 2–10% FBS, resuspended in 2% FBS PBS, filtered (Flowmi^®^ strainers, 40 *µ*m), and stained with DAPI immediately before sorting. Sorting was performed on a BD FACSAria (BD Diva software). Fluorescence-minus-one (FMO) controls were included in all experiments. Primary AML samples were sorted into CD34^+^ and CD34^−^ fractions; in cases with *>*50% CD34^+^ cells, the CD34^+^CD38^−*/*low^ population was also sorted. Sorted cells were collected in 50–100 *µ*l of ice-cold 2–10% FBS PBS or RPMI-1640. Post-sort viability was assessed using a LUNA dual fluorescence counter (Logos Biosystems). To enrich for immature populations that are often underrepresented in unsorted bone marrow mononuclear cells while preserving the broader cellular composition of each sample, the sorted fractions were recombined immediately prior to 10x Genomics library preparation. Specifically, CD34^+^ and CD34^-^ fractions (and CD34^+^CD38^-/low^ fractions when available) were pooled before loading onto the Chromium platform. The proportion of cells contributed by each fraction was recorded for each sample and is provided in(Supplementary Table 3).

### 4.3 Flow cytometry analysis of normal bone marrow and AML sampels

Normal bone marrow samples were stained with Fixable Viability Dye eFluor™ 455UV (Thermo Fisher Scientific, #65-0868-14) according to the manufacturer’s instructions, followed by FcR Blocking Reagent (Miltenyi Biotec, #130-059-901). Extracellular staining was performed using the following antibodies: anti-CD45 BUV615 (BD Biosciences, #751472), anti-CD34 APC (BioLegend, #343510), anti-CD90 PE (BioLegend, #328110), anti-CD90 BV510 (BioLegend, #328126), anti-CD38 PE–Texas Red (Thermo Fisher Scientific, #MHCD3817), anti-CD45RA RB670 (BD Biosciences, #571965), anti-CD123 BUV805 (BD Biosciences, #751840), anti-IL1RAP BV421 (BD Biosciences, #748107), anti-CD84 PE-Vio770 (Miltenyi Biotec, #130-119-038), anti-CD371 BUV661 (BD Biosciences, #750311), anti-CD3 PerCP (BioLegend, #344814), anti-CD14 BV786 (BD Biosciences, #741034), anti-CD19 BV650 (BioLe-gend, #363026), and anti-CD56 BUV563 (BD Biosciences, #612928). For antigen density quantification, anti-CD371 PE (BD Biosciences, #562566), anti-CD84 PE (Miltenyi Biotec, #130-119-035), and anti-IL1RAP PE (Bio-Techne, #FAB676P) were used for CD371, CD84, and IL1RAP quantification respectively, in conjunc-tion with the BD Quantibrite™ PE Phycoerythrin Fluorescence Quantitation Kit (BD Biosciences, #340495), across NBM and AML samples. All samples were acquired on a Cytek® Aurora spectral cytometer or a BD FACS Discover S8 cell sorter, following the manufacturer’s instructions. Healthy progenitor popula-tions were identified using established immunophenotypic gating strategies as fol-lows: haematopoietic stem cells (HSCs; CD34^+^CD38^−^CD90^+^CD45RA^−^), multipotent progenitors (MPPs; CD34^+^CD38^−^CD90^−^CD45RA^−^), lymphoid-primed multipotent progenitors (LMPPs; CD34^+^CD38^−^CD90^−^CD45RA^+^), common myeloid progeni-tors (CMPs; CD34^+^CD38^+^CD123^+^CD45RA^−^), granulocyte-monocyte progenitors (GMPs; CD34^+^CD38^+^CD123^+^CD45RA^+^), and megakaryocyte-erythrocyte progeni-tors (MEPs; CD34^+^CD38^+^CD123^−^CD45RA^−^). Mature populations were identified as CD14^+^ monocytes, CD3^+^ T cells, and CD19^+^ B cells. In AML samples, leukaemic blasts were analysed following exclusion of CD3^+^ and CD19^+^ cells.

### 4.4 Single-cell workflow, library preparation, and sequencing

Single-cell experiments were performed on the 10x Genomics Chromium platform using 5^*′*^ chemistry with Feature Barcode technology. Libraries were prepared following the manufacturer’s recommendations (Chromium Next GEM Single Cell 5^*′*^ v2 Dual Index, CG000330 Rev D). For each run, ∼20,000 cells were loaded (expected recovery ∼50%, yielding ∼10,000 cells).

Gel bead-in-emulsion (GEM) droplets were generated in the Chromium Controller and subjected to reverse transcription (45 min) before overnight storage at 4 °C. Subsequent cDNA amplification and clean-up followed 10x protocols. Paediatric AML and healthy control samples underwent 12 cycles of cDNA amplification. ADT libraries were amplified for 8 cycles; GEX libraries for 16 cycles (1–25 ng input) or 14 cycles (26–50 ng input).

ADT libraries were sequenced on a NextSeq2000 (Illumina) with dual indexing (26–10–10–90 cycles) at ∼5,000 reads/cell. GEX libraries (AML and healthy controls) were sequenced on a NovaSeq (Illumina) at ∼50,000 reads/cell (26–10–10–190 cycles).

### 4.5 Single-cell RNA sequencing analysis

Single-cell RNA-seq data were processed using Cell Ranger v7.1.0 for alignment and quantification against the human reference genome GRCh38 (refdata-gex-GRCh38-2020-A) (Supplementary Table 2). Ambient RNA and ADT counts were removed with CellBender[43] (false positive rate = 0.01 for RNA; 0.1 for ADT). ADT counts were further denoised using TotalVI from the scArches package[44]. Downstream analysis and quality control were performed in Scanpy.

Low-quality cells were filtered as follows. Each sample required a minimum of 10 genes per cell. For each cluster, a signature identity score was calculated using Scanpy’s rank genes groups function, defined as the number of distinct marker genes per cluster (adjusted *p <* 0.05, log fold change *>* 0.5, expressed in *>*10% of cells per group). Clusters with fewer than 300 genes were labelled low quality. After aggregating clustering results across all samples, a threshold of *<*600 detected genes and *<*800 total mRNA counts confidently removes all low-quality clusters while retaining high-identity clusters. Doublets were identified and removed using Scrublet with default parameters (expected_doublet_rate = 0.06; n_neighbors = round 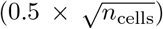)[45]. Cells with *>*20% mitochondrial gene expression were excluded. Counts were normalized using the delta method, and 3,000 highly variable genes were selected using the seurat v3 flavor (span = 0.3). Dimensionality reduction was performed with 50 principal components (PCs) computed using svd solver=“arpack”. A neighbor graph was built with 20 neighbors, 50 PCs, and cosine distance. UMAP was generated with min dist=0.3 and spread=1.2. Clustering was performed using the Leiden algorithm with resolution = 2.

### 4.6 Malignant and normal identification

Individual patient samples were integrated with healthy controls to create a joint dataset. Counts were normalized using the delta method, and 800 highly variable genes were selected using batch_key = sample and span = 0.3, followed by scaling with max value = 10. Principal component analysis (PCA) was computed using svd solver = “arpack”, and a K-nearest neighbors graph was constructed with 100 neighbors and 30 PCs. UMAP embeddings were generated with min_dist = 0.3 and spread = 1.2. Clustering was performed using the Leiden algorithm with a resolution of 2. For each Leiden cluster, an occupancy score was calculated as the fraction of cells from each patient relative to the total cells in that cluster. To refine cell malignancy classification, we applied an iterative KNN-based voting strategy. Initially, cells were labelled as malignant or normal based on sample type (“AML” → malignant; “Healthy” →normal). For each iteration, the proportion of healthy cells among the 50 nearest neighbors of each cell was computed. Cells with at least 20% healthy neighbors were reclassified as normal; otherwise, they retained their malignant label. This process was repeated until no further changes occurred or a maximum of 20 iterations was reached.

### 4.7 Fusion detection

Single-cell gene-fusion detection was performed via FusionInspector[46] in single-end mode, on Read2 .fastq files from 10x single cell sequencing experiments. Target fusions (Supplementary Table 3) were selected from previous studies[6, 47], including fusions from cytogenetic analysis on our case series; target fusions are included in canonical orientation (GeneA–GeneB) and in reversed orientation (GeneB–GeneA) to also investigate chimeric reads produced by reciprocal rearrangements. For downstream single-cell analyses, names of fusion-supporting reads from FusionInspector output have been used to extract cell barcodes and UMI from Read1 fastq files using custom scripts. Fusion Fragment Per Million (FFPM) values have been recalculated to take into account PCR duplicates, as follows: FFPM = (fusion supporting read counts / library size) *×* 1,000,000, where fusion_supporting_read_counts is the count of JunctionReads with unique barcode+UMI configuration, and library_size is the sum of gene counts in raw_feature_bc_matrix.h5 files from CellRanger analysis. Fusion-Inspector detected chimeric reads for 8 out of 9 expected fusions from cytogenetic analysis in 11 out of 12 fusion-positive samples (true positives); 6 of them have been detected only in true-positive samples. CBFB::MYH11 and KMT2A::AFDN show low FFPM levels of chimeric reads also in cytogenetic negative samples (false positives) attributable to background noise (Extended Data Fig. 3a); for each sample we calculated robust z-score as follows: *z*^∗^ = 0.6745 *×* (*x*_*i*_ ^−^ Median)*/*MAD, and true positive samples showed extremely high values (*z*^∗^ = 23.60–52.18) compared to false positives (*z*^∗^ = ^−^ 1.62–3.93), confirming the specificity of fusion detection. Notably, in AML10, KMT2A::AFDN shows high FFPM values (*z*^∗^ = 16.40–35.20), comparable to true positive samples; for this reason, we suspect AML10 carries an undetected KMT2A::AFDN fusion event, and we will consider it fusion-positive for downstream analyses. In 6 true positive samples, we also detected chimeric reads from the corresponding reciprocal fusion (Extended Data Fig. 3a). Reads from 7 unexpected additional fusions were detected (Extended Data Fig. 3a). TFG::ADGRG7 (and its reciprocal ADGRG7::TFG) is a previously reported polymorphic germline fusion, without clinical/pathological impact[48]. The other unexpected fusions show extremely low FFPM values compared to true positives (Welch *t*-test, *p* = 0.004) (Extended Data Fig. 3b), attributable to background noise.

### 4.8 InferCNV

To distinguish malignant from normal cells in each sample, copy number variation (CNV) was inferred from single-cell transcriptomic data using infercnvpy (Sturm, G., ICBI Lab, https://github.com/icbi-lab/infercnvpy), a scalable Python reimplementation of the Broad Institute’s InferCNV. The method computes per-cell log fold changes relative to a reference population by subtracting reference mean expression from each gene, smoothing values across a sliding genomic window, centering by cell median, and applying noise filtering to suppress low-magnitude variation. Healthy reference cells from our paediatric dataset were supplied as the normal reference population, and all remaining parameters were left at their defaults.

### 4.9 Annotation

Healthy bone marrow samples were annotated using the Zeng bone marrow reference[8]. Batch effects were corrected using the Harmony algorithm[49], and the first 30 Harmony-corrected principal components were used to construct the KNN graph and compute UMAP embeddings. AML patient cells were then projected onto this reference using the Symphony package[50].

### 4.10 Non-negative matrix factorization

Consensus non-negative matrix factorization (cNMF)[51] was applied to identify transcriptional programs within malignant cells. Raw count data from each sample were used as input. For each sample, highly variable genes (*n* = 2,000) excluding ribosomal genes were selected, and NMF was performed with 6 components and 200 iterations using the cNMF framework. Consensus programs were obtained using a density threshold of 0.1 to remove low-quality programs across patient cells. The resulting usage matrix and top 50 genes per program were used for downstream analysis. Python implementation of the AUCell method from decoupleR[52] was used to calculate LSC and blast score enrichment across cells using LSC and blast gene signatures [10–13, 16, 18]. To interpret biological relevance, program usage was correlated with known leukemia stem cell (LSC) and blast signatures. Programs with the highest average correlation to LSC signatures were annotated as LSC-high programs, and blast-associated programs were selected based on positive correlation with blast Score (Spearman correlation *>* 0.15) and low correlation with the LSC program. Pairwise Jaccard similarity was computed between all selected gene sets, defined as the size of the intersection divided by the size of the union of the sets. The resulting similarity matrix was converted into a distance matrix (1 − Jaccard similarity), and hierarchical clustering was performed using the average linkage method to visualize relationships between programs.

### 4.11 K-means clustering of programs

To identify shared cell populations across patients within the LSC–blast gene space, gene expression values from all selected programs were extracted and subjected to K-means clustering. To determine the optimal number of clusters, K-means was performed for a range of cluster numbers (*k* = 1–19), and the within-cluster sum of squares (WCSS) was computed for each *k*. The “elbow method” was used to select the point at which additional clusters no longer substantially reduced WCSS, indicating the optimal cluster number. Based on this analysis, seven clusters were chosen. K-means clustering was then applied to the selected gene expression data with *k* = 7, using the k-means++ initialization with 15 random starts (n init = 15).

### 4.12 ADT-QC, ECF calculation, and parameter optimisation

After denoising ADT counts using TotalVI from scArches and applying log normalization, a residual background signal remained. To define confidently detected ADTs, we examined correlations among all ADTs. While Isotype 1 and 3 showed little correlation with most ADTs, we observed that a high correlation persisted for one specific antibody even after background removal. To quantify the true signal relative to background, we calculated the Pearson correlation of each ADT with its corresponding RNA modality (representing real signal) and compared it to the ADT’s correlation with Isotype 2. ADTs that exhibited low correlation to RNA and high correlation to this isotype were subsequently removed. To define a threshold for ECF calculation (fraction of positive cells within each group), we modeled the distribution of counts across all cells with a two-component Gaussian mixture model and set the threshold as the midpoint between the component means, thereby distinguishing true signal from background. For quality control, we retained only cells with at least 14 detected ADTs and 200 counts per cell. ADT counts were then binarized using the determined threshold, classifying cells with counts above the cutoff as positive. Pareto analysis was performed using the Python paretoset package, with LSC ECF and HSC ECF as input variables.

### 4.13 External single-cell cohort analysis

FASTQ files from the external s ingle-cell cohort were processed using the same Cell Ranger version as our cohort (v7.1.0). QC and downstream analyses were performed in the same way as described for our cohort. A Random Forest classifier was trained using the Scikit-learn Python package on our cohort using malignant and normal cell annotations, and this model was applied to the external cohort to classify cells as malignant or normal. Model training included 5-fold cross-validation to ensure robust performance. For clustering, genes derived from our NMF analysis were used to cluster cells within the external cohort using K-means clustering to identify LSC-like populations.

### 4.14 scTranslator and scArches model

To predict the ADTs within the external single-cell cohort, we employed scTranslator, a pre-trained generative model that draws inspiration from natural language translation to facilitate the cross-modal prediction of protein abundance from RNA expression. Commencing with the pre-trained weights published by the original authors, we conducted nine distinct experiments. These trials involved three versions of the scTranslator model: 10k, 160k, and 2M, which had been pre-trained on datasets of progressively increasing scale (over 18,000 bulk samples; over 160,000 human PBMCs plus 18,000 bulk samples; and over 2 million human cells plus 18,000 bulk samples, respectively). Each model was then subjected to one of three second-stage fine-tuning strategies: utilising the original authors’ weights directly without further fine-tuning; fine-tuning for a single epoch; or fine-tuning for five epochs. This fine-tuning was performed using the paired RNA and ADT data from our own cohort. For each of the nine configurations, we evaluated performance by calculating the Jaccard similarity between the top 10 predicted protein markers and the top 100 RNA markers for every patient. Amongst the configurations tested, the scTranslator 2 M model, when fine-tuned for five epochs, yielded the greatest overlap. Consequently, this configuration was selected to predict the proteomic profiles for the external cohort.

To extend protein expression measurements to samples profiled by RNA-seq alone, we employed a transfer learning approach using totalVI implemented in the scArches package—a probabilistic deep generative model that jointly models RNA and protein expression within a variational autoencoder framework. A reference totalVI model was pre-trained on our CITE-seq cohort containing paired RNA and ADT measurements across multiple batches. For each RNA-only query dataset, raw count data were used as input, allowing query cells to be mapped onto the reference model’s latent space without requiring observed protein data. Query data were loaded into the pre-trained reference model using TOTALVI.load_query_data with frozen expression weights (freeze expression=True) to preserve the reference latent representation. The model was then fine-tuned on the query data for up to 200 epochs with no weight decay, enabling adaptation of batch-specific components while retaining the learned gene–protein relationships from the reference. Normalized protein expression was subsequently imputed by sampling from the posterior distribution (*n* = 25 samples) and averaging across all reference batches, yielding a batch-corrected imputed protein matrix for each query dataset.

To evaluate imputation accuracy, predictions were benchmarked against ground-truth ADT measurements from our CITE-seq cohort. For each patient sample, the top 10 most highly expressed ADT features were identified from both the reference and each prediction, forming per-sample ADT sets. These sets were compared using the Jaccard similarity index—defined as the size of the intersection divided by the size of the union of the two ADT sets—which captures a method’s ability to recover the correct protein identity per patient independently of expression scale. Jaccard scores were computed per sample and averaged across all patients to produce a single scalar similarity score per prediction method, enabling direct ranking and comparison of imputation approaches.

### 4.15 LSC clustering and projection

For in-depth analysis of the LSC-like population, the SAM[53] (Self-Assembling Manifold) algorithm was applied to this group using *k* = 30 and max_iter = 20, with normalization enabled, to identify genes with high manifold relevance, as previously described for identifying adult AML LSC groups[13]. The top 1,000 genes ranked by SAM weights were used to recompute RNA embeddings. Log-normalized counts were used as input, and PCA was performed on this gene subset, followed by batch effect correction using Harmony integration with sample identity as the batch key and 20 maximum iterations. A neighborhood graph was then constructed using the Harmony-corrected PCA space with cosine distance and a neighborhood size proportional to the dataset size. UMAP was computed on this graph to generate the RNA embeddings, which were subsequently used for Leiden clustering (resolution = 1) to obtain a finer granularity of subpopulations. LSC-like cells from Lasry et al.[28] were projected onto our LSC reference using a FAISS-based[54] nearest-neighbor search on the shared gene expression space. Cell type labels for query cells were assigned by transferring the most frequent label among their nearest reference neighbors identified through FAISS L2 similarity search. The projected UMAP coordinates for each query cell were then computed as the weighted average of its reference neighbors’ UMAP positions, preserving the reference embedding structure.

### 4.16 Inferring LSC group abundance in RNA-seq data

The abundance of each leukaemic stem cell (LSC) group is represented by a gene expression signature, comprising a linear equation with coefficients as signed by non-negative LASSO regression for each constituent gene. The gene signature was obtained using 50 marker genes from each LSC subtype. For each bulk RNA-seq sample, LASSO was applied by modeling the bulk expression profile as a linear combination of single-cell-derived LSC signatures. The regression coefficients we re co nstrained to be non-negative, reflecting that each LSC group contributes positively to the bulk signal. After fitting, coefficients for each sample were normalized to sum to one, yielding estimated proportions of each LSC group within that sample. These normalized values represent the relative abundance of LSC groups across patients. Gene expression signatures were derived from CPM-normalized single-cell RNA-seq data and applied to bulk RNA-seq data from both patient cohorts.

### 4.17 Survival analysis

Survival analyses were performed using the lifelines package using the ECF outcome. Differences b etween the two groups were assessed using the log-rank test, while comparisons across more than two groups were performed using a multivariate log-rank test. Kaplan–Meier estimators were fitted for each group, and survival curves were plotted with the function kmf.fit from this package.

## 5 Data availability

Processed single-cell RNA and surface protein (ADT) data generated in this study are available through the Synapse platform: https://www.synapse.org/Synapse:syn70497164.

## 6 Code availability

Custom scripts used for data processing and visualisation are available at https://github.com/sinakane/pAML or from the corresponding author upon request.

## 7 Author contributions

S.K. performed single-cell analyses and generated figures. M.O.E. conducted experiments and contributed to data analysis. I.M.M., T.X., E.K., C.D., C.R. and A.C. performed experiments. E.G.C., G.H., and H.M. provided input on data analysis. J.B. and R.T. provided clinical samples and patient data. E.G., S.C., and F.M. contributed to data analysis and figure preparation. A .G. and P.J.A. supervised the work. A.G. and S.K. wrote the manuscript with input from all authors. All authors reviewed and approved the final version.

## Acknowledgments

We thank the Human Technopole National Facilities and UCL facilities for their contribution to data generation and processing, in particular the National Facility for Genomics and the Human Technopole National Facility for Light Imaging, including the FLARE Flow Cytometry Unit, the UCL Great Ormond Street Institute of Child Health Flow Cytometry Facility and to the UCL Genomics Facility. Patient samples were provided by the VIVO Biobank and Great Ormond Street Hospital. This work was supported by the National Institute for Health and Care Research Great Ormond Street Hospital Biomedical Research Centre, Cancer Research UK Children and Young People Award (A.G.), the AIRC Start-Up Grant (A.G. ID 30423), the Leukaemia UK John Goldman Fellowship (A.G. 2018/JGF/003), and the Rosetrees Trust Fund (A.G., M700) and institutional funding from Human Technopole. The views expressed are those of the authors and not necessarily those of the National Health Service, the NIHR, or the Department of Health. S.K., A.C., and C.R. are members of the European School of Molecular Medicine (SEMM). M.O.E. was supported by a Children with Cancer UK Clinician PhD Fellowship.

## References

[1] Alexander, T. B. et al. Decreased relapse rate and treatment-related mortality contribute to improved outcomes for pediatric acute myeloid leukemia in successive clinical trials. Cancer 123, 3791–3798 (2017).

[2] Chen, J. & Glasser, C. L. New and emerging targeted therapies for pediatric acute myeloid leukemia (AML). Children 7, 12 (2020).

[3] Khoury, J. D., Solary, E., Abla, O. et al. The 5th edition of the World Health Organization Classification of Haematolymphoid Tumours: Myeloid and Histiocytic/Dendritic Neoplasms. Leukemia 36, 1703–1719 (2022).

[4] Arber, D. A., Orazi, A., Hasserjian, R. P. et al. International Consensus Classification of Myeloid Neoplasms and Acute Leukaemias: integrating morphologic, clinical, and genomic data. Blood 140, 1200–1228 (2022).

[5] Döhner, H. et al. Diagnosis and management of AML in adults: 2022 recommendations from an international expert panel on behalf of the ELN. Blood 140, 1345–1377 (2022).

[6] Bolouri, H. et al. The molecular landscape of pediatric acute myeloid leukemia reveals recurrent structural alterations and age-specific mutational interactions. Nat. Med. 24, 103–112 (2018).

[7] Umeda, M. et al. A new genomic framework to categorize pediatric acute myeloid leukemia. Nat. Genet. 56, 281–293 (2024).

[8] Zeng, A. G. X. et al. Single-cell transcriptional atlas of human hematopoiesis reveals genetic and hierarchy-based determinants of aberrant AML differentiation. Blood Cancer Discov. 6, 307–324 (2025).

[9] Lambo, S. et al. A longitudinal single-cell atlas of treatment response in pediatric AML. Cancer Cell 41, 2117–2135.e12 (2023).

[10] Eppert, K. et al. Stem cell gene expression programs influence clinical outcome in human leukemia. Nat. Med. 17, 1086–1093 (2011).

[11] Gentles, A. J., Plevritis, S. K., Majeti, R. & Alizadeh, A. A. Association of a leukemic stem cell gene expression signature with clinical outcomes in acute myeloid leukemia. JAMA 304, 2706–2715 (2010).

[12] Gal, H. et al. Gene expression profiles of AML-derived stem cells reveal similarity to hematopoietic stem cells. Leukemia 20, 2147–2154 (2006).

[13] Ng, S. W. K. et al. A 17-gene stemness score for rapid determination of risk in acute leukaemia. Nature 540, 433–437 (2016).

[14] Elsayed, A. H. et al. A six-gene leukemic stem cell score identifies high-risk pediatric acute myeloid leukemia. Leukemia 34, 735–745 (2020).

[15] Zhang, Y., Jiang, S., He, F. et al. Single-cell transcriptomics reveals multiple chemoresistant properties in leukemic stem and progenitor cells in pediatric AML. Genome Biol. 24, 199 (2023).

[16] Mumme, H. et al. Single-cell analysis reveals altered tumor microenvironments of relapse- and remission-associated pediatric acute myeloid leukemia. Nat. Commun. 14, 6209 (2023).

[17] Ohlstrom, D. et al. Single-cell analysis of pediatric acute myeloid leukemia samples uncovers treatment-resistant stem and mast cells. Leukemia 39, 2721–2734 (2025).

[18] Zeng, A. G. X. et al. A cellular hierarchy framework for understanding heterogeneity and predicting drug response in acute myeloid leukemia. Nat. Med. 28, 1212–1223 (2022).

[19] Pei, S. et al. A novel type of monocytic leukemia stem cell revealed by the clinical use of venetoclax-based therapy. Cancer Discov. 13, 2032–2049 (2023).

[20] Sango, J. et al. RAS-mutant leukaemia stem cells drive clinical resistance to venetoclax. Nature 636, 241–250 (2024).

[21] Bhagwat, A. S. et al. Cytokine-mediated CAR T therapy resistance in AML. Nat. Med. 30, 3697–3708 (2024).

[22] Haubner, S., Subklewe, M. & Sadelain, M. Honing CAR T cells to tackle acute myeloid leukemia. Blood 145, 1113–1125 (2025).

[23] Haubner, S. et al. Cooperative CAR targeting to selectively eliminate AML and minimize escape. Cancer Cell 41, 1871–1891.e6 (2023).

[24] Geyer, M. B. et al. CD371-targeted CAR T cells secreting interleukin-18 exhibit robust expansion and clear refractory acute myeloid leukemia. Blood 146, 3163– 3174 (2025).

[25] Gottschlich, A. et al. Single-cell transcriptomic atlas-guided development of CAR-T cells for the treatment of acute myeloid leukemia. Nat. Biotechnol. 41, 1618– 1632 (2023).

[26] Perna, F. et al. Integrating proteomics and transcriptomics for systematic combinatorial chimeric antigen receptor therapy of AML. Cancer Cell 32, 506–519.e5 (2017).

[27] Zhang, C. et al. Filtering cells with high mitochondrial content depletes viable metabolically altered malignant cell populations in cancer single-cell studies. Genome Biol. 26, 91 (2025).

[28] Lasry, A. et al. An inflammatory state remodels the immune microenvironment and improves risk stratification in acute myeloid leukemia. Nat. Cancer 4, 27–42 (2023).

[29] Kwon, J. et al. Single-cell mapping of combinatorial target antigens for CAR switches using logic gates. Nat. Biotechnol. 41, 1593–1605 (2023).

[30] Zhang, X. et al. An immunophenotype-coupled transcriptomic atlas of human hematopoietic progenitors. Nat. Immunol. 25, 703–715 (2024).

[31] Tabula Sapiens Consortium et al. The Tabula Sapiens: A multiple-organ, single-cell transcriptomic atlas of humans. Science 376, eabl4896 (2022).

[32] Uhlén, M. et al. Tissue-based map of the human proteome. Science 347, 1260419 (2015).

[33] Liu, L. et al. A pre-trained large generative model for translating single-cell transcriptomes to proteomes. Nat. Biomed. Eng. (2025).

[34] Trad, R. et al. Chimeric antigen receptor T-cells targeting IL-1RAP: a promising new cellular immunotherapy to treat acute myeloid leukemia. J. Immunother. Cancer 10, e004222 (2022).

[35] Zhu, Y. et al. Identification of CD84 as a potent survival factor in acute myeloid leukemia. J. Clin. Invest. 135, e176818 (2025).

[36] Zhang, H. et al. Characteristics of anti-CLL1-based CAR-T therapy for children with relapsed or refractory acute myeloid leukemia: a multi-center efficacy and safety interim analysis. Leukemia 36, 2596–2604 (2022).

[37] Pigazzi, M. et al. CD84 is a specific target for acute myeloid leukemia CAR-T cell therapy. Nat. Commun. (2026).

[38] Pérez-Amill, L. et al. A novel chimeric antigen receptor T-cell therapy targeting CD84 for the treatment of acute myeloid and T-cell lymphoblastic leukemias. Leukemia 39, 2432–2441 (2025).

[39] Jin, X. et al. First-in-human phase I study of CLL-1 CAR-T cells in adults with relapsed/refractory acute myeloid leukemia. J. Hematol. Oncol. 15, 88 (2022).

[40] Pei, K. et al. Anti-CLL1-based CAR T cells with 4-1BB or CD28/CD27 stimulatory domains in treating childhood refractory or relapsed acute myeloid leukemia. Cancer Med. 12, 9655–9661 (2023).

[41] Wang, H. et al. The tandem CD33–CLL1 CAR-T as an approach to treat acute myeloid leukemia. Blood Transfus. 23, 338–347 (2025).

[42] Xie, D., Jin, X., Sun, R. et al. Bicistronic CAR-T cells targeting CD123 and CLL1 for AML to reduce the risk of antigen escape. Transl. Oncol. 34, 101695 (2023).

[43] Fleming, S. J. et al. Unsupervised removal of systematic background noise from droplet-based single-cell experiments using CellBender. Nat. Methods 20, 1323– 1335 (2023).

[44] Lotfollahi, M. et al. Mapping single-cell data to reference atlases by transfer learning. Nat. Biotechnol. 40, 121–130 (2022).

[45] Wolock, S. L., Lopez, R. & Klein, A. M. Scrublet: computational identification of cell doublets in single-cell transcriptomic data. Cell Syst. 8, 281–291.e9 (2019).

[46] Haas, B. J. et al. Targeted in silico characterization of fusion transcripts in tumor and normal tissues via FusionInspector. Cell Rep. Methods 3, 100467 (2023).

[47] Shiba, N. et al. Transcriptome analysis offers a comprehensive illustration of the genetic background of pediatric acute myeloid leukemia. Blood Adv. 3, 3157–3169 (2019).

[48] Chase, A. et al. TFG, a target of chromosome translocations in lymphoma and soft tissue tumors, fuses to GPR128 in healthy individuals. Haematologica 95, 20–26 (2010).

[49] Korsunsky, I. et al. Fast, sensitive, and accurate integration of single-cell data with Harmony. Nat. Methods 16, 1289–1296 (2019).

[50] Kang, J. B. et al. Efficient and precise single-cell reference atlas mapping with Symphony. Nat. Commun. 12, 5890 (2021).

[51] Kotliar, D. et al. Identifying gene expression programs of cell-type identity and cellular activity with single-cell RNA-seq. eLife 8, e43803 (2019).

[52] Badia-i Mompel, P. et al. decoupleR: ensemble of computational methods to infer biological activities from omics data. Bioinformatics Adv. 2, vbac016 (2022).

[53] Tarashansky, A. J., Xue, Y., Li, P., Quake, S. R. & Wang, B. Self-assembling manifolds in single-cell RNA sequencing data. eLife 8, e48994 (2019).

[54] Johnson, J., Douze, M. & Jégou, H. Billion-scale similarity search with GPUs. arXiv (2017).

